# Loss of function of metabolic traits in typhoidal *Salmonella* without apparent genome degradation

**DOI:** 10.1101/2024.02.14.580360

**Authors:** Leopoldo F. M. Machado, Jorge E. Galán

## Abstract

*Salmonella enterica* serovar Typhi and Paratyphi A are the cause of typhoid and paratyphoid fever in humans, which are systemic life-threatening illnesses. Both serovars are exclusively adapted to the human host, where they can cause life-long persistent infection. A distinct feature of these serovars is the presence of a relatively high number of degraded coding sequences coding for metabolic pathways, most likely a consequence of their adaptation to a single host. As a result of convergent evolution, these serovars shared many of the degraded coding sequences although often affecting different genes in the same metabolic pathway. However, there are several coding sequences that appear intact in one serovar while clearly degraded in the other, suggesting differences in their metabolic capabilities. Here, we examined the functionality of metabolic pathways that appear intact in *S*. Typhi but that show clear signs of degradation in *S*. Paratyphi A. We found that, in all cases, the existence of single amino acid substitutions in *S.* Typhi metabolic enzymes, transporters, or transcription regulators resulted in the inactivation of these metabolic pathways. Thus, the inability of *S*. Typhi to metabolize Glucose-6-Phosphate or 3-phosphoglyceric acid is due to the silencing of the expression of the genes encoding the transporters for these compounds due to point mutations in the transcriptional regulatory proteins. In contrast, its inability to utilize glucarate or galactarate is due to the presence of point mutations in the transporter and enzymes necessary for the metabolism of these sugars. These studies provide additional support for the concept of adaptive convergent evolution of these two human-adapted *Salmonella enterica* serovars and highlight a limitation of bioinformatic approaches to predict metabolic capabilities.

## Introduction

*Salmonella enterica* continues to be a major public health challenge (1–4). Based on their antigenic composition, *Salmonella enterica* is classified in multiple serovars (5), which exhibit different host specificity and pathogenicity. In addition, according to their pathogenesis, *Salmonella enterica* serovars are often grouped into those that cause systemic infection (i.e. typhoidal serovars) and those that cause self-limiting gastroenteritis (i.e. non-typhoidal serovars) (6–9). *Salmonella* Typhi and *Salmonella* Paratyphi A are the most common typhoidal serovars that affect humans, where they cause typhoid and paratyphoid fever, which affect approximately 20 million people every year resulting in approximately 150,000 deaths world-wide (3, 6, 10–12). A subset of individuals that recover from these illnesses harbor these organisms in the gall bladder for life, where they serve as reservoirs for future infections.

*Salmonella* Typhi and *Salmonella* Paratyphi A arose independently through evolution from their non-typhoidal common ancestors (13, 14). However, up to a quarter of their genomic material was subsequently exchanged through recombination (15). The process of their adaptation to a single host combined with their unique lifestyle are partially reflected by the presence of a significantly higher number of degraded coding sequences or pseudogenes coding for virulence factors (e. g. type III secretion effectors) or various metabolic pathways (e. g. central anaerobic metabolism) when compared to broad host *S. enterica* serovars (13–15). Importantly, genome degradation is observed in areas of the chromosome that were not subject to exchange, thus exhibiting evidence of adaptive convergent evolution. Comparison of degraded coding sequences in different serovars can be a useful tool to gain insight into specific adaptations to a given niche. In fact, this type of analysis has allowed the identification of metabolic signatures specifically associated with serovars that cause gastroenteritis or systemic infection (16, 17).

*S*. Typhi and *S*. Paratyphi A exhibit significant differences in the set of degraded coding sequences, in particular, several genes that encode metabolic pathways (13, 14). This is surprising because these organisms share the same niche (i. e. the human host) and exhibit indistinguishable clinical presentation and pathogenic features (i. e. both serovars cause typhoid fever and systemic and persistent infection) (18). Here we have examined coding sequences that are degraded in one typhoidal serovar (i.e. *S*. Paratyphi A) but that are apparently intact in another (i.e. *S*. Typhi). We found that in the cases examined, specific point mutations in the *S*. Typhi metabolic genes or their regulators have rendered them non-functional. These results further support the concept of adaptive convergent evolution shaping the metabolic pathways of typhoidal *Salmonella*. In addition, these findings uncover a limitation of comparative genomic approaches to detect phenotypic differences between related bacteria.

## Results

### *S.* Typhi and *S.* Paratyphi A show differences in degraded coding sequences for metabolic pathways

Based on previously published comparisons between typhoidal and non-typhoidal *Salmonellae* (13, 14, 16, 17), and using the *S.* Typhimurium LT2 strain as reference, we compiled a list of genes encoding well-characterized or putative metabolic pathways that are either absent or annotated as pseudogenes in *S*. Typhi (strains CT18 and Ty2) or Paratyphi A (strains ATCC 9150 and AKU12601) (Table S1). As previously noted (13–15, 19), and in line with the concept of convergent evolution during adaptation to extraintestinal infection in the human host, there is significant overlap in the metabolic pathways that have been silenced in both organisms. However, this inactivation often occurs through the degradation of different genes within the same pathway. Intriguingly and as noted before (13), there are instances of metabolic pathways lost in one serovar but not in the other, where the coding genes appear to be intact. For example, this is the case for the glucose-6-phosphate, 3-phosphoglyceric acid, and glucarate/galactarate utilization pathways in *S*. Typhi, which have been degraded in *S.* Paratyphi A, or the galactitol and L-rhamnose utilization pathways in *S*. Paratyphi A, which have been degraded in *S.* Typhi (Table S1). This observation is intriguing as it suggests potential metabolic differences in these bacterial pathogens. In turn, these differences may reflect partially distinct environments during their pathogenic cycles, which could have potentially influenced the selection of different metabolic capabilities. This would be surprising considering their shared mechanisms of pathogenesis, clinical manifestations, and the ability to cause persistent infection. On the other hand, this observation raises the question whether these apparently intact pathways are truly operational. To address this issue, we investigated the functionality of a set of metabolic pathways where the coding genes are degraded in *S.* Paratyphi A but are putatively intact in *S*. Typhi.

### Glucose-6-phophate (G6P) sensing and transcription regulation are impaired in *S*. Typhi despite the absence of gene degradation

Glucose-6-phosphate (G6P) is abundantly present within the cytosol of eukaryotic host cells (20), although it is absent within the *Salmonella*-containing vacuole (SCV) (21). As some *S. enterica* serovars such as *S*. Typhimurium can gain access to the cytosolic environment (22), the observation of potential differences in the ability to utilize G6P by *S*. Typhi and *S*. Paratyphi A is of potential physiological significance. The import of G6P is mediated by the UhpT antiporter, whose expression is strictly controlled by an unconventional two-component regulatory system composed of the membrane-localized UhpC sensor and UhpB kinase/phosphatase, and the cytoplasmic response regulator UhpA (23, 24). In this system, UhpC directly senses G6P triggering a conformational change in the interacting kinase UhpB, which in turn phosphorylates and activates the response regulator UhpA. Phosphorylation increases the affinity of UhpA for the *uhpT* promoter (P*uhpT*) thus initiating transcription (Fig. 1A). Although the *uhpA*, *uhpB*, *uhpC*, and *uhpT* coding sequences are apparently intact in *S*. Typhi, they are degraded in *S*. Paratyphi A (Table S1).

**Fig. 1.**
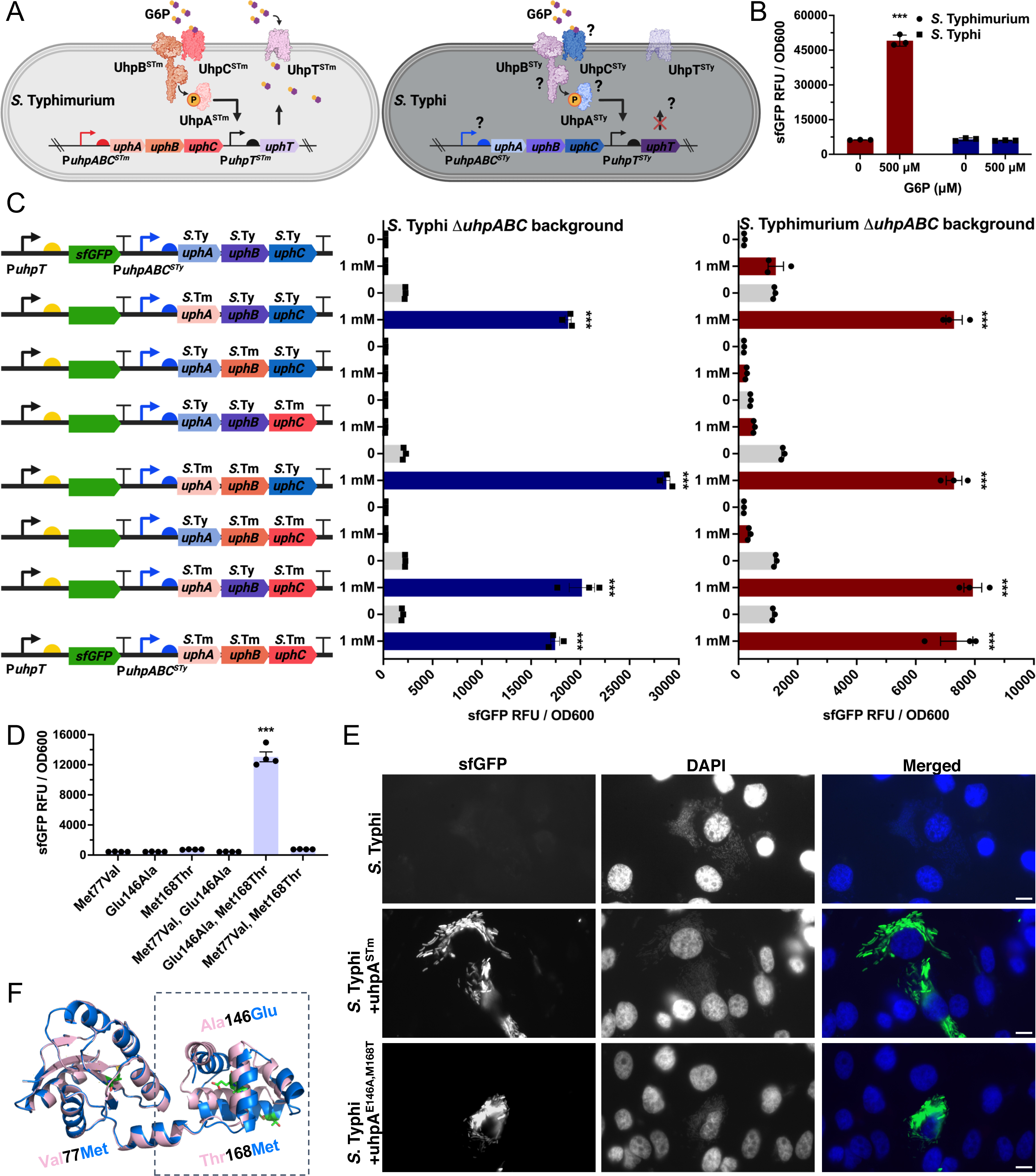
Glucose-6-phosphate sensing and transcriptional regulation are impaired in *Salmonella* Typhi. (**A**) Diagram of the glucose-6-phosphate (G6P) sensing, transcription regulation, and transport in *Salmonella enterica*. G6P (purple hexagon) is imported into the bacterial cell by the UhpT antiporter, whose expression is strictly controlled by an unconventional two-component regulatory system composed of the membrane-associated UhpC sensor and UhpB kinase/phosphatase, and the cytoplasmic response regulator UhpA. In this system, UhpC directly senses G6P triggering a conformational change in the interacting kinase UhpB, which in turn phosphorylates and activates the response regulator UhpA. Phosphorylation increases the affinity of UhpA for the *uhpT* promoter (P*uhpT*) thus initiating transcription. The potential steps where the sensing, regulation and transport could be impaired in *S*. Typhi are noted. (**B**) Transcriptional response of the *S*. Typhimurium and *S*. Typhi G6P biosensors after growth in LB medium containing G6P (500 µM). The sfGFP fluorescence signal for *S*. Typhimurium (circle on red bar) or *S*. Typhi (square on blue bar) in the presence or absence of G6P is shown. (**C**) Investigation of the functionality of UhpA, UhpB, and UhpC regulatory proteins. Each *S*. Typhi gene was swapped for its *S*. Typhimurium homolog in the plasmid backbone containing the G6P biosensor as indicated in the diagram. The resulting plasmids were then introduced into the *S*. Typhi or *S.* Typhimurium *ΔuhpABC* mutant backgrounds, and the resulting strains were grown in M9 medium in the presence or absence of G6P (1mM). (**D**) Investigation of the functionality of each of the different *uhpA* mutations present in *S*. Typhi in comparison of its functional homolog in *S*. Typhimurium. Plasmids encoding the G6P biosensor and expressing *S*. Typhi UhpA mutants in which residues had been individually changed to the amino acid present in its *S*. Typhimurium homolog as indicated (i. e. UhpA^Met77Val^, UhpA^Glu146Ala^, or UhpA^Met168Thr^ ) were introduced into the *S.* Typhi Δ*uhpABC* mutant background. The resulting strains were grown into M9 medium in the presence of G6P (1mM). In (**B**-**D**) the sfGFP fluorescence signal normalized to cell density (sfGFP RFU / OD600) was measured. Values indicate the mean ± SEM of n= 3 or 4 replicates per condition. Asterisks denote statistically significant differences relative to the corresponding uninduced sample determined using Anova with Tukey’s multiple comparisons test ***p < 0.001, (**B** and **D**), or unpaired two-tailed t-test ****p < 0.0001 (**C**). (**E**) Functionality of different UhpA variants expressed in *S*. Typhi in the context of cultured epithelial cell infection. HeLa cells were infected with wild-type *S*. Typhi or derivative strains expressing the *uhpA* allele from S. Typhimurium (*uhpA^STm^*) or an *S*. Typhi allele, which had been rendered functional by introducing changes based on the *S*. Typhimurium sequence (i. e. *uhpA^Glu146Ala,Met168Thr^*). Hela cells were infected with the different *S.* Typhi strains for 6 hours, fixed, stained with DAPI, and examined under a fluorescence microscope. In all the images, brightness and contrast were optimized for each of the individual color channels to maximize visual clarity. Scale bars, 10 μm. (**F**) Alignment of the modeled atomic structures of *S*. Typhi UhpA (light blue) and S. Typhimurium UhpA (light pink) showing structural variations in the C-terminal portion at the helix-turn-helix motif (boxed). The models were predicted using AlphaFold2 and aligned with Pymol. The amino acid differences between the two structures (Val77Met, Ala146Glu, and Thr168Met) are shown in green.

To investigate whether the G6P pathway was operational in *S*. Typhi we constructed a G6P biosensor/transcriptional reporter in which the P*uhpT* promoter drives the expression of super-folder GFP (sfGFP), as previously reported in *S*. Typhimurium (25–27) (Fig. S1A). We found that, as previously reported, the *S*. Typhimurium biosensor produced a robust fluorescence signal when the bacteria were grown in the presence of G6P. However, no signal was detected when *S*. Typhi encoding an equivalent biosensor was grown in the same media (Fig. 1B and S1B). Consistent with these observations, a bright fluorescence signal was detected in cytosolic *S*. Typhimurium after infection of HeLa cells, but no fluorescence signal was detected in cytosolic *S*. Typhi encoding an equivalent G6P biosensor (Fig. S1C). These results imply that despite the presence of apparently intact coding sequences, G6P sensing and/or transcriptional regulation is impaired in *S*. Typhi.

Comparison of the nucleotide sequence of the *uhp* loci in *S*. Typhi and *S*. Typhimurium detected an 80 bp insertion on the *S*. Typhi P*uhpABC* promoter as well as point mutations on the *uhpA*, *uhpB*, and *uhpC* coding sequences, which could be responsible for the observed differences in the expression of the G6P biosensor. We first investigated whether the 80 bp insertion in the *S*. Typhi P*uhpABC* promoter was responsible for the observed differences in the expression of G6P biosensor. To this aim, we swapped the P*uhpABC* promoters in the *S*. Typhi and *S*. Typhimurium biosensors and quantified their expression in the presence of G6P by flow cytometry. We found robust expression of the G6P biosensor driven by the P*uhpABC S*. Typhi promoter in *S*. Typhimurium but we observed no expression of the biosensor driven by the P*uhpABC S*. Typhimurium promoter in *S*. Typhi (Fig. S2A). Similar results were obtained with equivalent transcriptional reporters driving the expression of nanoluciferase (Fig. S2 B and C). These results indicated differences in the promoter sequences could not explain the lack of expression of the G6P biosensor in *S*. Typhi.

We then examined whether point mutations in the *S*. Typhi UhpC sensor, its interacting kinase UhpB, or the response regulator UhpA were responsible for the inability of *S*. Typhi to respond to the presence of G6P. We constructed transcriptional reporters in which the *S*. Typhi *uhpA*, *uhpB*, or *uhpC* coding sequences were alternatively replaced with those from *S*. Typhimurium and expressed them in the *S*. Typhi and S. Typhimurium *ΔuhpABC* mutant backgrounds. We found that all combinations expressing the UhpA response-regulator from *S.* Typhi were unable to activate the transcriptional reporter in the presence of G6P. In addition, expression in *S.* Typhi or S. Typhimurium of *uhpA* from *S.* Typhimurium in the context of *uhpB* and *uhpC* from S. Typhi led to robust expression of the transcriptional reporter in the presence of G6P (Fig. 1C). These results indicated that the inability of the G6P transcriptional reporter to operate in *S*. Typhi is at least in part due to mutations within the response regulator UhpA.

In comparison to its *S.* Typhimurium homolog, *S*. Typhi UhpA (UhpA^STy^) exhibits three amino acid changes: Val77Met, Ala146Glu, and Thr168Met (Fig. S3). We changed each one of the *S*. Typhi UhpA amino acids to the *S*. Typhimurium residues in the equivalent positions and examined the expression of the G6P transcriptional reporter. We found that simultaneously introducing in *S.* Typhi the *uhpA^Glu146Ala^* and *uhpA^Met168Thr^* mutations rescued the expression of the G6P transcriptional reporter indicating that these point mutations inactivated UhpA in *S*. Typhi (Fig. 1D). We also found that this *S*. Typhi mutant strain as well as an *S*. Typhi strain encoding the *S.* Typhimurium *uhpA* allele showed expression of the transcriptional reporter when gaining access to the cytosol of infected HeLa cells (Fig. 1E). These observations indicate that the mutations present in the *S.* Typhi *uhpA* allele have rendered this response regulator inactive. Alignment of the structures of UhpA^STy^ (blue at Fig 1F) and UhpA^STm^ (pink at Fig 1F) as predicted by AlphaFold2 revealed that the two mutations that led to the loss of function of UhpA in *S*. Typhi reside within its helix-turn-helix DNA-binding domain, indicating that, most likely, these point mutations impaired the ability of UhpA^STy^ to activate the P*uhpT* promoter. Taken together, our results indicate that two amino acid substitutions within the response regulator UhpA have rendered it inactive in *S*. Typhi thus impairing G6P sensing and transcriptional regulation of the *uhp* genes.

### Glucose-6-Phosphate transport is impaired in *S*. Typhi due to a loss-of-function mutation in UhpT

Since the transcriptional regulation of the G6P utilization genes is impaired in *S*. Typhi due to mutations in the response regulator UhpA, it is predicted that the metabolism of this sugar would be impaired in *S*. Typhi. Consistent with this prediction and in contrast to *S*. Typhimurium, *S*. Typhi was unable to grow in M9 minimal medium containing G6P as the sole carbon source. Intriguingly, *S*. Typhi expressing the *S*. Typhimurium UhpA response regulator was also unable to grow in minimal medium with G6P (Fig. 2A) suggesting that defects other than transcription regulation must also be contributing to the inability of *S*. Typhi to utilize this sugar. We therefore swapped the gene encoding the UhpT transporter for its counterpart from *S*. Typhimurium in the context of a *S*. Typhi strain that expressed the UhpA response regulator from *S*. Typhimurium. We found that the resulting mutant strain was able to robustly grow in M9 medium supplemented with G6P as the only carbon source (Fig. 2B). In comparison to *S*. Typhimurium, *S*. Typhi UhpT exhibits four amino acid substitutions: Asp18Asn, Pro27Ser, Gly61Arg, and Ala395Asp (Fig. S4). To investigate which of the mutations resulted in the loss of function, we changed each one of the *S*. Typhi amino acids to those present in its *S*. Typhimurium homolog and expressed the resulting constructs in the context of a *S*. Typhi strain encoding the *S*. Typhimurium UhpA response regulator (which in *S*. Typhi carries a loss of function mutation, see above). We found that introducing in *S*. Typhi the *uhpT^Asp395Ala^* allele rescued the ability of *S*. Typhi to grow in M9 minimal medium supplemented with G6P as the only carbon source (Fig. 2C). Interestingly, the structure of UhpT predicted by AlphaFold2 indicates that alanine^395^ resides within an alpha helix that makes up the lumen of the predicted channel of the UhpT transporter (Fig 2D). Therefore, mutation of this residue most likely impairs the transport function of UhpT. Taken together, these results indicate that the ability to utilize G6P by *S*. Typhi is not only impaired by a mutation in the response regulator that controls expression of the transporter, but also by a mutation in the transporter itself that impairs its function.

**Fig. 2.**
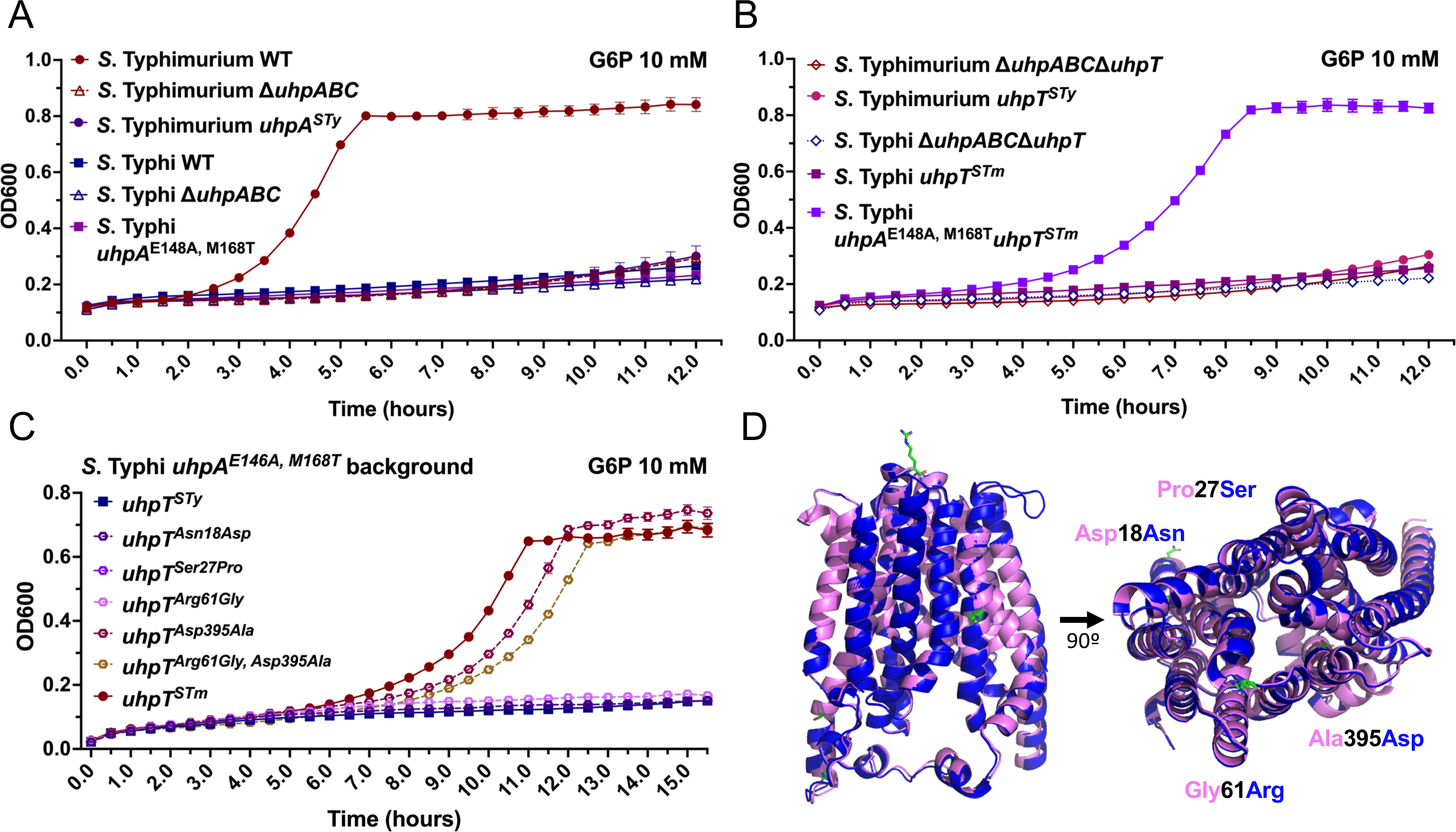
Glucose-6-Phosphate transport is impaired in *S*. Typhi due to a loss-of-function mutation in UhpT. (**A**-**C**) Growth kinetics of wild-type (WT) *S*. Typhimurium and *S*. Typhi or the indicated mutant strains. Bacteria were grown in M9 medium containing 10 mM G6P as the only carbon source. The OD600 values measured at 30 min intervals are shown. Values indicate the mean ± SEM of n=3 replicates per condition. (**D**) Alignment of the modeled atomic structures of *S*. Typhi (dark blue) and *S*. Typhimurium (pink) UhpT predicted with AlphaFold2 and aligned with Pymol. The amino acids that differ between the two structures (Asp18Asn, Pro27Ser, Gly61Arg, and Ala395Asp) are shown in green.

G6P has been hypothesized to serve as an important carbon source for *S*. Typhimurium growth within the cytosol of infected cells. Since *S*. Typhi is also able to gain access to the cytosol, we investigated whether the inability to utilize G6P could affect its ability to grow within this compartment. To address this issue, we made use the *S*. Typhi strain encoding UhpA and UhpT from S. Typhimurium, which can utilize G6P. Since growth within the cytosolic compartment is much faster than within the SCV, bacteria that can utilize the metabolites available in the cytosol may be able to grow better in this compartment, particularly at later times after infection. We found no difference in the total number of CFU of wild-type *S.* Typhi (unable to utilize G6P) with that of the strain expressing the *S*. Typhimurium UhpA and UhpT alleles, which can utilize this sugar (Fig. S5). Although this experiment did not directly measure growth in the cytosol, given that in HeLa cells late in infection most of the CFU are generated in the cell cytosol (replication in the vacuole is substantially slower), these results argue that the inability to use G6P does not seem to substantially handicap the ability of *S*. Typhi to grow within the cytosol of cultured epithelial cells.

### A loss of function mutation in the PgtA response regulator impairs 3-phosphoglyceric acid metabolism in *S*. Typhi

The metabolite 3-phosphoglyceric acid (3PG) is an important intermediate in the aerobic glycolysis pathway in eukaryotic cells (28). Importantly, stimulation of innate immune receptors in macrophages by bacterial ligands promotes the switch of glucose metabolism from oxidative phosphorylation to aerobic glycolysis (i.e. the "Warburg effect") with accumulation of the glycolytic intermediate 3PG (29, 30). It has been recently reported that 3PG is essential for *S*. Typhimurium intracellular replication and systemic virulence (31). Consequently, potential differences in the ability to utilize 3PG by the typhoidal serovars *S*. Typhi and *S*. Paratyphi A could have important implications for their pathogenesis. The transport of 3PG in *S. enterica* is mediated by the PgtP transporter, whose expression is controlled by the unconventional two-component regulatory system composed of the membrane-localized PgtC sensor and PgtB kinase/phosphatase, and the cytoplasmic PgtA response regulator (32, 33) (Fig. 3A). While the *pgtB* coding sequence is clearly degraded in *S.* Paratyphi A, all the genes required for 3PG utilization are apparently intact in *S*. Typhi (Table S1).

**Fig 3.**
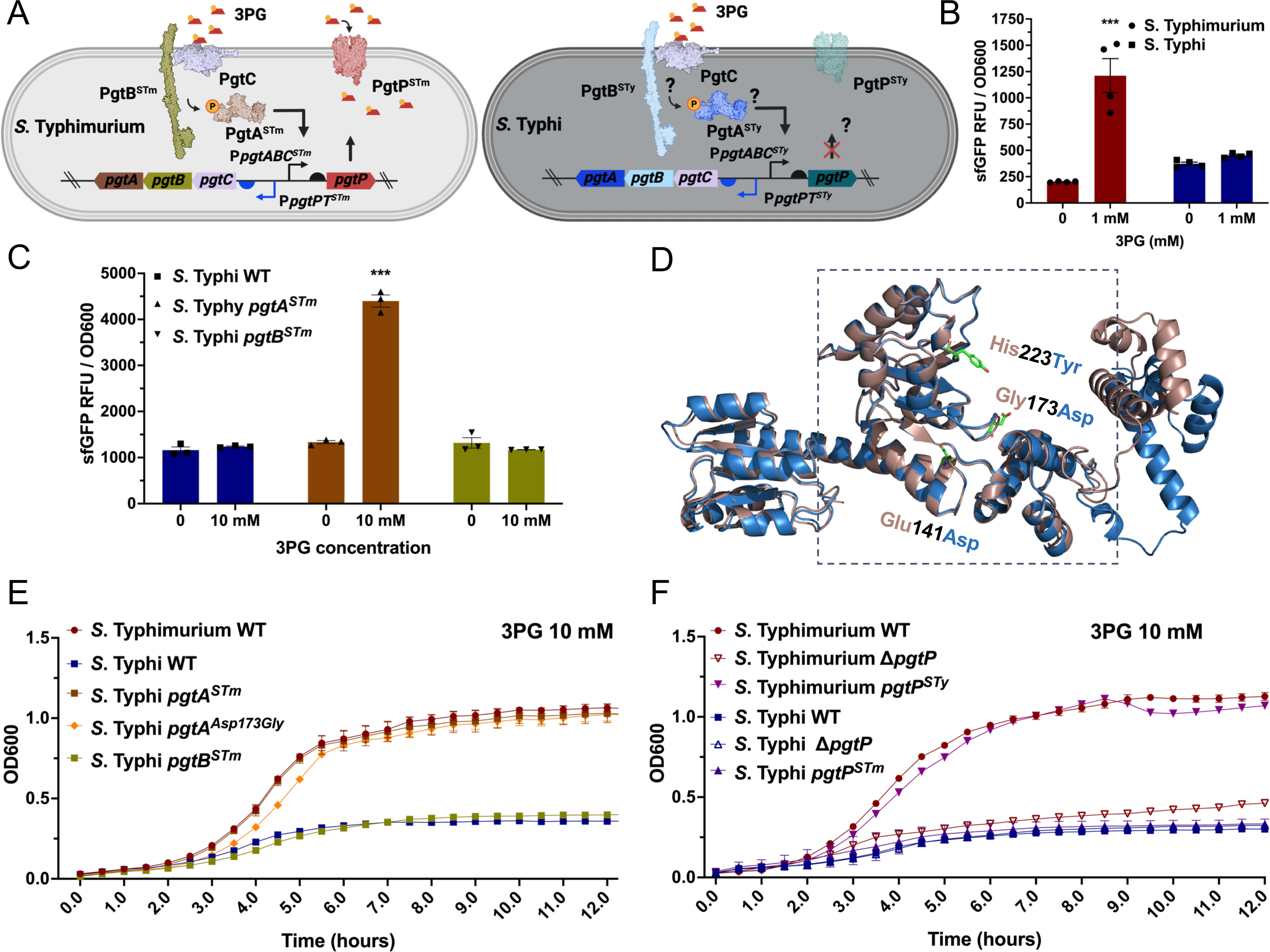
The metabolism of 3-phosphoglycerate is impaired in *Salmonella* Typhi due to a loss of function mutation in the PgtA response regulator. (**A**) Diagram of 3PG sensing, transcription regulation, and transport in *Salmonella enterica*. PgtC senses 3PG (red trapezoid) triggering its interaction with the PgtB histidine kinase/phosphatase that phosphorylates the PgtA response regulator. Phosphorylated PgtA, in turn, activates the P*gtpP* promoter leading to the expression of the PgtP transporter. The potential steps where the sensing, regulation and transport could be impaired in *S*. Typhi are noted. (**B**) Transcriptional response of the *S*. Typhimurium and *S*. Typhi strains encoding the 3PG biosensors after growth in LB medium containing 3PG (1 mM). The sfGFP fluorescence signal for *S*. Typhimurium (circle on red bar) or *S*. Typhi (square on blue bar) in the presence or absence of 3PG is shown. Values indicate the mean ± SEM of n=3 replicates per condition. (**C**) Investigation of the functionality of the S. Typhi PgtA and PgtB regulatory proteins. The indicated *S*. Typhi genes were swapped for their *S*. Typhimurium homologs and the resulting mutant strains were tested for their ability to sense 3PG with the P*pgtP-sfGFP* biosensor. Bacteria were grown in LB medium for 20 h in the presence or absence of 3PG (1mM), and the fluorescence signal (normalized to cell density) was measured (sfGFP RFU/OD600) for each strain. Values in (**B**) and (**C**) are the mean ± SEM of n=3-4 replicates per condition. Asterisks denote statistically significant differences relative to the corresponding uninduced sample determined using Anova with Tukey’s multiple comparisons test. ***p < 0.001. (**D**) Alignment of the modeled atomic structures of *S*. Typhi (blue) and *S*. Typhimurium (pink) PgtA predicted with AlphaFold2 and aligned with Pymol. The amino acids that differ between the two structures (Glu141Asp, Gly173Asp, and His223Tyr) are shown in green. (**E** and **F**) Growth kinetics of wild type (WT) *S*. Typhimurium and *S*. Typhi or the indicated mutant strains. Bacteria were grown in M9 + 0.05% casamino acids medium containing 10 mM 3PG as the only carbon source. The OD600 values measured at 30 min intervals are shown. Values indicate the mean ± SEM of n=3 replicates per condition.

To investigate if the 3PG pathway is functional in *S*. Typhi, we designed a 3PG biosensor in which the *S.* Typhi P*pgtP* promoter drives the expression of sfGFP on a plasmid system. We then introduced the plasmid into both *S.* Typhimurium and *S*. Typhi. We observed that while *S*. Typhimurium harboring the 3PG biosensor produced a high fluorescence signal when the bacteria were grown in the presence of 3PG, no signal was detected when *S*. Typhi harboring the same plasmid was grown under identical conditions (Fig. 3B). These results indicate that, while the *S*. Typhi P*pgtP* promoter is functional (there are only 5 nucleotide differences between *S*. Typhi and *S.* Typhimurium in the 435 bp region that contains the P*pgtABC* and P*pgtP* promoters), the sensing and/or transcriptional regulation of the 3PG utilization genes are impaired in *S.* Typhi.

We sought to identify which components of the 3PG sensing and metabolism were impaired in *S*. Typhi (Fig 3A). Compared to its *S*. Typhimurium homologs, there are 3 amino acid substitutions in the *S.* Typhi PgtB kinase and the PgtA response regulator, while the PgtC sensor is identical in both serovars. Thus, we investigated whether the point mutations in PgtA or PgtB were the cause of the absence of signal for *S*. Typhi in the presence of 3PG. We constructed versions of the 3PG biosensor where the *S*. Typhi *pgtA* and *pgtB* coding sequences were replaced with those from *S*. Typhimurium and expressed them in a *S*. Typhi *ΔpgtAB* mutant strain. Expression in *S.* Typhi of *pgtA* from *S.* Typhimurium in the context of *pgtB* and *pgtC* from *S*. Typhi led to robust expression of the transcriptional reporter in the presence of 3PG (Fig. 3C and Fig. S6), confirming that mutations in *pgtA* impair the transcription regulation of 3PG utilization genes in *S.* Typhi.

In comparison to *S*. Typhimurium, the response regulator PgtA in *S*. Typhi (PgtA^STy^) exhibits three amino acid substitutions: Glu141Asp, Gly173Asp, and His223Tyr (Fig 3D and Fig. S7). We changed each one of these amino acids in *S*. Typhi for those present in *S*. Typhimurium and tested the expression of the 3PG transcriptional reporter. We observed that the introduction of the *pgtA^Asp173Gly^* mutation rescued the expression of the 3PG transcriptional reporter indicating that this point mutation inactivated PgtA in *S*. Typhi (Fig S8A and S8B). The AlphaFold2 model of PgtA indicates that Gly173 is located within a domain that is predicted to interact with σ54 suggesting that mutations in this residue may prevent the interaction of the *S.* Typhi response regulator with the sigma factor thus precluding expression of the 3PG utilization genes (Fig. 3D).

In the case of the glucose-6-phosphate, the inability of *S*. Typhi to utilize this sugar is due not only to defects in sugar sensing and transcription regulation but also to mutations in the transporter itself. We therefore tested whether 3PG transport was impaired in *S*. Typhi by examining the growth of the *S*. Typhi *pgtA^STm^* or the *S*. Typhi *pgtA^Asp173Gly^* mutants in M9 containing 3PG as the only carbon source (Fig 3E). Both strains were able to grow in this medium in a manner that was indistinguishable from *S*. Typhimurium wild type or an *S*. Typhimurium mutant expressing the *S*. Typhi 3PG transporter PgtP (Fig. 3F). Collectively, these results indicate that the inability of *S*. Typhi to utilize 3PG is due to the presence of a loss-of-function mutation in the response regulator PgtA.

### *S*. Typhi is unable to metabolize glucarate and galactarate due to loss-of-function mutations in the GudT transporter and the GudD and GarD dehydratases

In *S.* Paratyphi A the genes involved in glucarate and galactarate utilization, *gudT*, *ygcY*, *gudD* and *garD*, show clear signs of degradation and therefore are considered to be pseudogenes. In contrast, these coding sequences remain apparently intact in *S*. Typhi (Table S1). This observation is intriguing since the utilization of these sugars has been proposed to be only important for the growth of *Salmonella enterica* in the inflamed gut (34), which would not be relevant for *S.* Typhi, an extraintestinal pathogen. Therefore, we investigated whether these *S*. Typhi genes encode functional proteins. Glucarate and galactarate in *Salmonella enterica* is transported by the permease GudT and subsequently dehydrated to 5-dehydro-4-deoxy-D-glucarate by the dehydratases GudD and GarD, respectively (35, 36). Subsequently, 5-dehydro-4-deoxy-D-glucarate is metabolized into D-glycerate by the aldolase GarL and the reductase GarR, thus feeding the central metabolism. The expression of these metabolic genes is controlled by the transcription factor CdaR, which senses these sugars in the cytosol and activates the promoters for the genes responsible for their uptake and metabolism (Fig 4A). To study the sensing and metabolism of glucarate and galactarate in *S*. Typhi, we first constructed biosensors consisting of the *S.* Typhi promoters for *gudTYD* (P*gudTYD*), *gardD* (P*garD*) or *garLRK* (P*garLRK*) driving the expression of sfGFP (Fig. S9A). We found that when these reporters were introduced into *S*. Typhimurium, they readily responded to the presence of glucarate and galactarate. In particular, the P*garLRK*-sfGFP reporter showed the most robust response. However, this reporter did not respond to the presence of glucarate or galactarate in *S*. Typhi (Figure S9B). This observation indicated that although the promoters for these genes seem to be functional, either their transcriptional control by the CdaR regulator, or the uptake of these sugars by the GudT transporter must be defective in *S*. Typhi. In addition to glucarate and galactarate, CdaR also responds to the presence of glycerate, which unlike the sugars, does not require specific transport as it diffuses through the membrane (35). Therefore, we tested the response of the P*garLRK*-sfGFP reporter when *S*. Typhi was grown in the presence of glycerate (Fig 4B). We found that under these growth conditions, the reporter displayed a robust response indicating that CdaR is functional in *S.* Typhi, and therefore the inability of this reporter strain to respond to glucarate and galactarate must be due to defects in the transport of these sugars through GudT.

**Fig 4.**
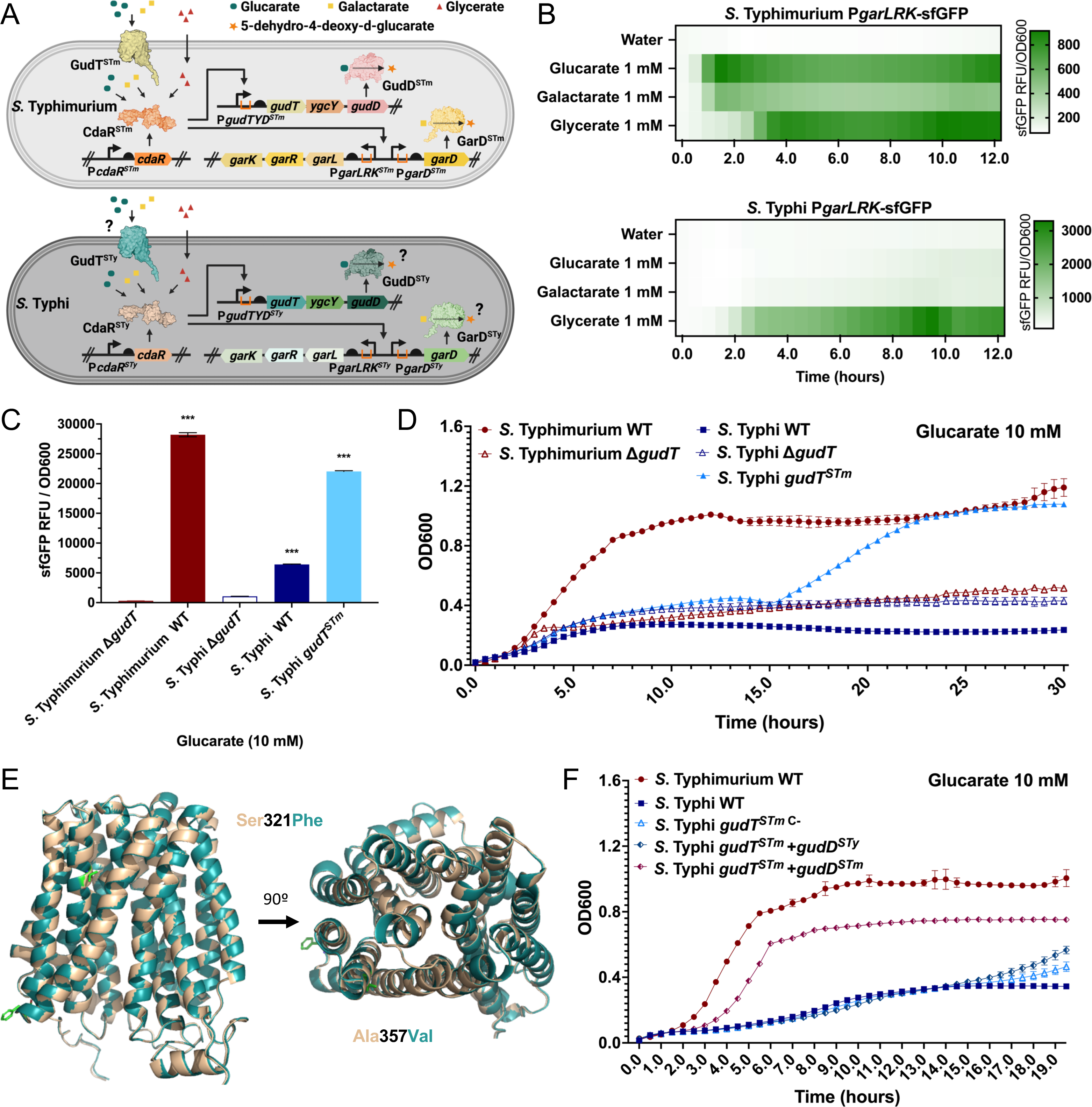
The glucarate and galactarate metabolism are impaired in *Salmonella* Typhi due to loss of function mutations in the GudT transporter and the GudD dehydratase. (**A**) Diagram of glucarate and galactarate sensing, transcription regulation, transport, and metabolism in *Salmonella enterica*. Glucarate (green circles) and galactarate (yellow squares) are transported into the bacterial cell by the permease GudT and converted to 5-dehydro-4-deoxy-d-glucarate (orange star) by the dehydratases GudD and GarD, respectively. 5-dehydro-4-deoxy-d-glucarate is further metabolized into D-glycerate (red triangles) by the aldolase GarL and the reductase GarR. The expression of these genes is controlled by the transcription factor CdaR, which senses glucarate, galactarate and D-glycerate in the cytosol and activates the P*gudTYD*, P*garD* and P*garLRK* promoters. The potential steps where the sensing, regulation, transport, or metabolism could be impaired in *S*. Typhi are noted. (**B** and **C**) Wild-type *S*. Typhimurium and S. Typhi or the indicated mutant strains all harboring the PgarL_*sfGFP* transcriptional reporter were grown in LB medium containing glucarate, galactarate or glycerate (as indicated) at 10 mM (or water as a negative control) and the fluorescence signal (normalized to cell density) was measured (sfGFP RFU/OD600) for each strain. Values indicate the mean ± SEM of n= 3 replicates per condition. Asterisks denote statistically significant differences determined using Anova with Tukey’s multiple comparisons test. ***p < 0.001. (**D**) Growth kinetics of wild-type *S.* Typhimurium and *S.* Typhi or their isogenic mutants *ΔgudT, ΔgudT* and *gudT* as indicated. Growth was monitored in M9 + 0.05% casamino acids medium containing 10 mM glucarate as the only carbon source. The OD600 measured at 30 min intervals are shown and values represent the mean ± SEM of n=3-4 replicates per condition. (**E**) Alignment of the modeled atomic structures of *S*. Typhi (Teal) and *S*. Typhimurium (Beige) GudT predicted with AlphaFold2 and aligned with Pymol. The amino acids that differ between the two structures (Ser321Phe and Ala357Val) are shown in green. (**F**) Growth kinetics of *S.* Typhi *gudT^STm^* expressing plasmid-borne *gudD*^STm^ and *gudD*^STy^, as indicated. Growth was monitored in M9 medium + 0.05% casamino acids containing 10 mM glucarate as the only carbon source. The OD600 measured at 30 min intervals are shown and values represent the mean ± SEM of n=3-4 replicates per condition.

To test this hypothesis, we constructed a strain of *S.* Typhi expressing the GudT homolog from *S*. Typhimurium and examined the response of the P*garLRK*-sfGFP transcriptional reporter to the presence of glucarate or galactarate. In contrast to wild type or the *ΔgudT* mutant strains, the *S*. Typhi strain expressing *S*. Typhimurium GudT showed a robust response of the P*garLRK*-sfGFP transcriptional reporter in the presence of these sugars (Fig. 4C; Fig. S9C). The *S*. Typhi strain expressing the GudT transporter from *S*. Typhimurium was able to grow in M9 minimal medium containing glucarate or galactarate as the sole source of carbon. However, in comparison to *S*. Typhimurium, growth in galactarate was limited and growth in glucarate was preceded by an extended lag period (Fig. 4D and Fig. S9D). These results indicated that the glucarate and galactarate GudT transporter is defective in *S*. Typhi. Comparison of the amino acid sequence of GudT in *S*. Typhi and *S*. Typhimurium showed the presence of two amino acid changes at positions Ser321 (*S*. Typhi GudT^Ser321Phe^) and Ala357 (*S*. Typhi GudT^Ala357Val^) (Fig. S10). The structure of the GudT transporter predicted by AlphaFold2 showed that Ala^357^ is located on an alpha helix that forms the putative channel of the GudT transporter (Fig. 4E). It is therefore possible that the presence of a valine at this position may disrupt the GudT channel activity in *S.* Typhi.

The observation that, relative to *S*. Typhimurium the *S*. Typhi strain expressing the GudT transporter from *S*. Typhimurium showed an extended lag period in media with galactarate or glucarate as the only carbon source raised the issue as to whether other elements of the glucarate/galactarate metabolism pathway may also been defective in *S*. Typhi. After its transport through the GudT transporter, glucarate and galactarate are metabolized by the GudD and GarD dehydratases, respectively. When compared to *S*. Typhimurium homologs, the amino acid sequences of the *S*. Typhi GudD and GarD exhibit several amino acid substitutions that could account for the observed deficiencies in the ability of *S*. Typhi to utilize these sugars: Ile43Val, Gly78Asp and Ala159Val in GudD, and Ser237Asn in GarD (Fig S11). To explore this possibility, we expressed *S*. Typhimurium GudD or GarD in a *S*. Typhi strain expressing the GudT transporter from *S*. Typhimurium. We found that the resulting strains were able to grow in M9 minimal media supplemented with glucarate or galactarate, respectively, as the only carbon source to levels equivalent to those of *S*. Typhimurium, although a lag period was also apparent (Fig. 4F and S12).

Taken together these results indicate that *S*. Typhi is unable to utilize galactarate and glutarate due to mutations in the GudT transporter, as well as in GudD and GarD, which are critical enzymes for the metabolism of these sugars. Overall, these results provide further demonstration that the metabolic changes that occurred in *S.* Typhi presumably through the process of adaptation to a single host and a systemic type of infection included not only gene degradation but more subtle, specific changes that has led to the inactivation of critical metabolic genes.

## Discussion

The process of adaptation to persistent infection in a single host has resulted in a significant accumulation of degraded coding sequences or pseudogenes in the typhoidal *Salmonella enterica* serovars *S*. Typhi and *S*. Paratyphi A (13, 14). While on average broad-host non-typhoidal *Salmonella* serovars exhibit ∼1 % of degraded coding sequences, typhoidal Salmonellae carry significant higher number of pseudogenes (>4%). Comparison of the degraded coding sequences in *Salmonella* serovars that cause systemic or gastrointestinal infection has been useful to glean specific metabolic traits necessary for replication in these two different environments (17). This comparison has also revealed several coding sequences that although clearly degraded in *S*. Paratyphi A, appear to be intact in *S*. Typhi (13). Many of these genes encode important metabolic pathways, which could suggest potential differences in the host niches occupied by these two typhoidal *Salmonella enterica* serovars. Given the high similarity between the pathogenesis of these two human-adapted typhoidal serovars, this observation was surprising.

To gain insight into this issue, we closely examined the functionality of a subset of *S.* Typhi genes that encode functions necessary for the utilization of key carbon sources: glucose-6-phosphate, which is abundant within the cytosol of mammalian cells (20); 3-phosphoglyceric acid, a product of aerobic glycolysis that accumulates in activated macrophages and has been shown to be a key signal to reprogram the metabolism of *S*. Typhimurium within macrophages (31); and glucarate and galactarate, which have been shown to be critical for the growth of *S*. Typhimurium in the inflamed gut (34). The genes related to the utilization of these carbon sources appear intact in *S*. Typhi, with none or very limited number of substitutions when their sequences are compared to the *S*. Typhimurium homologs. We found that, in all cases, while all the promoters of the transcriptional units are intact, the metabolic pathways encoded by these genes are inactive due to point mutations in one or more of the regulatory components, transporters, or metabolic enzymes. These mutations are highly conserved across the more than 1,000 *S*. Typhi isolates whose genome sequences are currently available in the NCBI data bases. Thus, the inability to utilize G6P is due to two amino acid substitution in the DNA-binding domain of the response regulator UhpA and a single amino acid substitution in the G6P transporter UhpT. Similarly, the inability to metabolize 3PG is due to a single amino acid substitution in the response regulator PgtA although the PgtP transporter retains its ability to function at wild-type levels, both when expressed in *S*. Typhi or *S*. Typhimurium. Therefore, the inability to utilize both G6P and 3PG by *S*. Typhi is due to the silencing of the expression of the genes encoding the necessary transporters due to point mutations in the regulatory components. In contrast, the presence of point mutations in the transporter and enzymes necessary for their metabolism are responsible for the inability of *S*. Typhi to utilize glucarate and galactarate. It is noteworthy that in this case, *S*. Typhi is still able to express these inactive enzymes and transporter. It is thought that genes that are either not expressed or that encode inactive products are eventually degraded due to the accumulation of mutations. This appears to be the case for the genes that encode the G6P, 3PG, and glucarate/galactarate utilization pathway in *S*. Paratyphi A. However, the observation that the homologous sequences remained intact in *S*. Typhi or that in one instance (i. e. the glucarate utilization pathway) are still expressed despite the presence of loss-of-function mutations suggest that the inactivation of these pathways must have been a relatively recent event.

Although our analyses involved only a subset of genes that are degraded in *S*. Paratyphi A but not in *S.* Typhi, it is likely that a similar situation may apply to other metabolic pathways that are degraded in one serovar but not in the other. For example, *sty3536*, which codes for a putative tartrate transporter, which is critical for tartrate utilization, is clearly degraded in *S.* Paratyphi A but appears intact in *S*. Typhi. However, both *S*. Typhi and *S*. Paratyphi A are unable to metabolize tartaric acid (Fig. S13). Comparison of the amino acid sequence of *S*. Typhi STY3536 with its homolog in *S*. Typhimurium, which can utilize tartaric acid, reveals the presence of one amino acid substitution that may lead to loss of function (Fig. S14). Furthermore, amino acid differences between the *S*. Typhi and *S*. Typhimurium homologs are also present within STY3537 and STY3538, which encode putative transcription regulators of this metabolic pathway (Fig. S14B and C). Similarly, genes coding for regulators, transporters and enzymes important for the metabolism of aspartate are clearly degraded in *S*. Paratyphi A but remain putatively intact in *S*. Typhi. For example, *dcuS*, which encodes the sensor-kinase of the two-component regulatory system that regulates the DcuABC-mediated uptake of aspartate (37), has point mutations in *S*. Typhi compared to its homolog in *S*. Typhimurium (Fig. S15A). Amino acid differences between the *S*. Typhi and *S*. Typhimurium homologs are also present on the aspartate chemo-attractant *tar* and on the enzyme AsnB (Fig. S15B and C) (38). Point mutations in these proteins may lead to loss of function and could explain *S*. Typhi’s inability to utilize aspartate. Another example is *hutU,* which encodes a hydratase that is essential for histidine utilization (39). This gene is degraded in *S.* Paratyphi A but appears intact in *S*. Typhi. However, *S*. Typhi was not able to utilize histidine, suggesting that the presence of single amino acid substitutions in *S*. Typhi HutU and/or the transcription regulator HutC relative to functional homologs are likely to be responsible for its inability to metabolize this amino acid (Fig. S16).

Our analyses confirm the remarkable convergence of the metabolism of the typhoidal serovar *S*. Typhi and *S*. Paratyphi A during the process of adaptation to the human host, suggesting a strong selection for the elimination of specific metabolic pathways. Interestingly, these metabolic pathways also appear degraded in other host adapted serovars such as *S*. Cholerasuis, *S*. Gallinarum, and *S.* Dublin (Table S1). It is possible that absence of these metabolites in the specific environments of the specific hosts where these pathogens may reside during persistent infection may provide the selection pressure to result in the loss of these metabolic capabilities. However, this remarkable convergence may indicate that the utilization of these carbon sources may somehow be deleterious for some not understood aspect of their physiology when in their specific hosts. More experiments will be required to test this hypothesis. Finally, our analyses also highlight a limitation in the prediction of metabolic capabilities based only on bioinformatic analyses.

## Materials and Methods

### Bacteria strains and cell lines

The *Salmonella* strains used in the experiments conducted in this study were derived from the *Salmonella* enterica serovar Typhimurium strain SL1344 (40), and *Salmonella* enterica serovar Typhi strain ISP2825 (41) and Salmonella enterica serovar Paratyphi A strain 3343 A-5 (Roy Curtiss III, strain collection) listed in Table S3. Mutants were constructed using standard recombinant DNA and allelic exchange using R6K suicide plasmids(42) in *E. coli* ß2163 Δ*nic35* as donor strain (43). Strains were routinely cultured in LB broth at 37 °C. The experiments using cultured cells were conducted using the HeLa human epithelial cell line. The cells were cultured in Dulbecco’s Modified Eagle Medium (DMEM, Gibco) supplemented with 10% Bovine Calf Serum (BCS) at 37 °C with 5% CO2 in a humidified incubator.

### Plasmid construction

All the plasmids used in this study were constructed using Gibson assembly cloning (44) and are listed in Table S2. Plasmids used for the biosensors/transcriptional reporters, the expression of the point mutants in *uhpA, uhpT and pgtA*, and the wild type *gudD* and *garD* enzymes were constructed using the medium copy p15A ori vector pAJM.657(45). The plasmids used for generation of the *Salmonella* chromosomal mutants were constructed using the R6K suicide plasmid pSB890 as described(46) and are listed in Table S2.

The G6P biosensor was constructed by amplifying the P*uhpT* promoter (-151 to -25 bp upstream to the *uhpT* coding sequence) from *S.* Typhi ISP2825, placing it upstream to the phage G10 RBS (47) and *sfGFP* in the p15A ORI pAJM.657 backbone generating the vector pSB6424. Derivatives of this plasmid encoding *uhpA uhpB and uhpC* from either *S*. Typhi or *S*. Typhimurium were constructed by amplifying these genes and the native promoters from the respective strains and subsequently cloning them into pSB6424. Derivatives of these plasmids encoding point mutations in the *S*. Typhi UhpA to residues present in the *S*. Typhimurium homolog (i. e. UhpA^Met77Val^, UhpA^Glu146Ala^, or UhpA^Met168Thr^) were constructed by site directed mutagenesis using standard molecular biology techniques. Plasmids expressing UhpT from *S*. Typhi or *S*. Typhimurium from their native promoters were constructed in the plasmid vector backbone pAJM.657 (45). Mutations in the *S*. Typhi UhpT to residues present in the *S.* Typhimurium UhpT homolog (i. e. UhpT^Asp18Asn^, UhpT^Ser27Pro^, UhpT^Arg61Gly^, or UhpT^Asp395Ala^) were generated by site directed mutagenesis following standard molecular biology techniques.

The 3PG biosensor was constructed by amplifying the P*pgtP* promoter (-435 to -25 bp upstream to the *pgtP* coding sequence) from *S*. Typhi ISP2825, placing it upstream to the Phage G10 RBS and *sfGFP* in the p15A ORI pAJM.657 backbone generating the vector pSB6546. Derivatives of this plasmid encoding PgtA and PgtB from either *S*. Typhi or *S*. Typhimurium were constructed by amplifying these genes and the native promoters from the respective strains and subsequently cloning them into pSB6546. Derivatives of these plasmids encoding point mutations in the S. Typhi PgtA to residues present in the S. Typhimurium homolog (i. e. PgtA^Asp141Glu^, PgtA^Asp173Gly^ and PgtA^Tyr223His^) were constructed by site directed mutagenesis using standard molecular biology techniques.

The glucarate/galactarate biosensors used in this study were constructed by amplifying the P*gudTYD*, P*garD* and P*garLRK* promoters (-520 bp upstream to the coding sequence of each operon) from the *S*. Typhi ISP2825 genome and cloning them upstream to a *sfGFP* gene into the pAJM.657 p15A ORI backbone, generating the vectors pSB6407, pSB6408, and pSB6409, respectively. To improve the signal levels, the PgarLRK^STy^-*sfGFP* biosensor was modified by replacing the native RBS with the phage G10 RBS, generating the glucarate/galactarate biosensor P*garLRK.*G10-s*fGFP* vector pSB6534. For the expression of *gudD,* its coding sequences from *S*. Typhi or *S*. Typhimurium were cloned in the pAJM.657 backbone downstream from the P*gudTYD* promoter consisting of 520 nucleotides upstream of the *gudT* plus 36 nucleotides of *gudT*’s actual coding sequence. For the expression of *garD,* its coding and promoter sequences (i. e. 520 bp upstream of its initiation codon) from *S*. Typhi or *S*. Typhimurium were cloned in the pAJM.657 backbone.

### Transcriptional reporter assays

Wild-type and mutant strains of *S*. Typhimurium and *S*. Typhi harboring the different transcriptional reporters were grown at 37 °C in LB supplemented with glucose-6-phosphate (G6P) (Roche) (concentrations ranging from 7.6 μM to 10 mM, as indicated), D-3-phosphoglyceric acid (3PG) (Sigma) (concentrations ranging from 1 mM to 10 mM, as indicated), glucaric acid (Sigma) (concentrations ranging from 1 mM to 10 mM, as indicated), or galactaric acid (Sigma) (concentrations ranging from 1 mM to 10 mM, as indicated) for at least 12 hours. Incubation and agitation were carried out made in final 200 μl volume in flat-bottom black wall 96 well plates (Costar) using a Spark multimode microplate reader (Tecan). The growth and sfGFP fluorescence signal were monitored with measurements at 30 min intervals (excitation wavelength of 485 nm, bandwidth 15 nm; emission wavelength of 530 nm, bandwidth of 15 nm). The background fluorescence of the medium without bacteria was subtracted from all readings, which were then normalized by the bacterial OD_600_ growth to give the final sfGFP RFU/OD_600_ values. Alternatively, *S.* Typhimurium and *S.* Typhi and indicated mutants harboring the P*uhpABC^STm^*-*NLuc* or P*uhpABC^STy^*-*NLuc* transcriptional were grown for 20 hours with agitation at 37°C, diluted 10 x into sterile water and processed with the Nano-Glo® Luciferase Assay System (Promega). The bioluminescence was measured using a Spark multimode microplate reader (Tecan).

### Analytical Flow Cytometry

Wild-type *S*. Typhimurium and *S*. Typhi, as well as the indicated mutant derivatives harboring the G6P biosensor P*uhpT*-*sfGFP* were grown in the presence of G6P as described above. The cultures were then washed two times and re-suspended in PBS to a final concentration of ∼10^6^ bacteria/ml and the fluorescence intensity measured at 488 nm excitation, 530/33 emission was analyzed for 10,000 bacteria using a BD Accuri C6 flow cytometer.

### Bacterial growth in minimal medium

Wild-type and mutant strains of *S*. Typhimurium and *S*. Typhi were grown in M9 minimal medium (M9 salts 6 g Na_2_HPO_4_, 3 g KH_2_PO_4_, 0.5 g NaCl, 1 g NH_4_Cl per liter; 2mM MgSO_4_; 0.1 mM CaCl_2;_ 50 μg/ml L-tryptophan; 50 μg/ml L-cysteine hydrochloride; 50 μg/ml L-histidine) supplemented with 10 mM G6P, 10 mM 3PG, 10 mM glucarate or 10 mM galactarate as the only carbon source, as indicated. Except when strains were grown in the presence of G6P, the media were supplemented with casamino acids 0.05% (w/v). In all cases, the *Salmonella* strains were grown in LB medium overnight at 37°C until stationary phase. Cultures were washed two times in PBS and diluted to an OD_600_ of 0.025 in M9 minimal medium containing one of the specific carbon sources, and incubated on 96 well plates (Costar) at 37°C with orbital agitation for at least 12 hours using the Spark multimode microplate reader (Tecan). Culture growth was monitored with measurements at 30 min intervals. The background of the medium without bacteria was subtracted from all readings to give the final OD_600_ values.

### Salmonella infections

For cultured HeLa cell infections, the different *Salmonella* strains were grown in a modified LB broth containing 0.3 M NaCl to induce the expression of the SPI-1 T3SS. Overnight cultures were diluted 1: 20 and grown to an OD_600_ of 0.9 (48). HeLa cells at a confluency of 80% were infected for 1 h with *S*. Typhimurium or *S*. Typhi in Hank’s balanced salt solution (HBSS) at multiplicities of infection (MOI) of 25 and 50, respectively. The infected cells were washed three times with PBS and incubated in DMEM containing 100 μg/ml gentamicin to kill extracellular bacteria. After 1 h, cells were washed once with PBS and further incubated in DMEM containing 10 μg/ml gentamicin for the remaining time of infection, as indicated in the figure legends. At the indicated times, cultures were lysed in 0.5 ml of PBS containing 0.2 % Na-deoxycholate and the number of colony forming units (CFU) was determined by dilution plating in LB agar plates. Bacterial intracellular growth was determined by dividing the CFU at 8 h by the CFU at 2 h after infection.

### Fluorescence Microscopy

HeLa cells grown on glass coverslips were infected with *S*. Typhimurium or *S*. Typhi carrying a plasmid with the G6P biosensor/transcriptional reporter P*uhpT*-*sfGFP* as indicated above. At 6h and 20 h post-infection, the samples were washed three times with PBS, fixed in 4% paraformaldehyde (PFA) for 15 min and washed with PBS again. The cells were then permeabilized with a buffer containing PBS, 3% BSA and 0.3% Triton-X, stained with DAPI (Sigma) for 5 min and washed three times with PBS. Finally, the slides were mounted using ProLong antifade Mountaunt (Thermo) and left to dry for 24 hs. Slides were imaged using an Eclipse TE2000 inverted microscope (Nikon) with an Andor Zyla 5.5 sCMOS camera controlled by the Micromanager software (https://www. micro-manager.org). Brightness and contrast were optimized for each of the individual color channels to maximize visual clarity.

### Genomic analysis

Degraded coding sequences (pseudogenes) for metabolic genes in *S.* Paratyphi A (strains ATCC 9150 and AKU_12601) that are putatively intact in *S.* Typhi (strains CT18 and Ty2) compared to the sequences in the reference *S.* Typhimurium strain LT2 were listed at Table S1 based on previously reported genomic comparisons(13, 14, 16, 17).

### Protein amino acid sequences alignments

Protein amino acid sequences were retrieved from the reference genomes of the serovars *S.* Typhi CT18 (AL513382.1), *S.* Paratyphi A ATCC 9150 (CP000026.1), *S.* Typhimurium LT2 (AE006468.2), *S.* Enteritidis P125109 (AM933172.1) and *S.* Agona SL483 (CP001138.1). Sequences were aligned using Clustal Omega (EMBL-EBI) and the alignment figures show the relevant regions containing the amino acid point mutations in *S.* Typhi.

### Protein structural models

Protein structures were modeled using the AlphaFold2 notebook from ColabFold executing the default parameters (49). Visualization and structural alignment were carried out using PyMol (50).

### Statistical analysis

Statistical significance was calculated either by Anova with Tukey’s multiple comparisons test or Student’s unpaired two-tailed t-tests. For most experiments, values indicate the mean ± SEM with n= 3-4 replicates per condition. Further details are provided in the figure legends.

## Acknowledgments

This work was supported by NIH Grant R01AI114618 to J.E.G.

## Supplementary Information

### Supplementary figures

**Figure S1.**
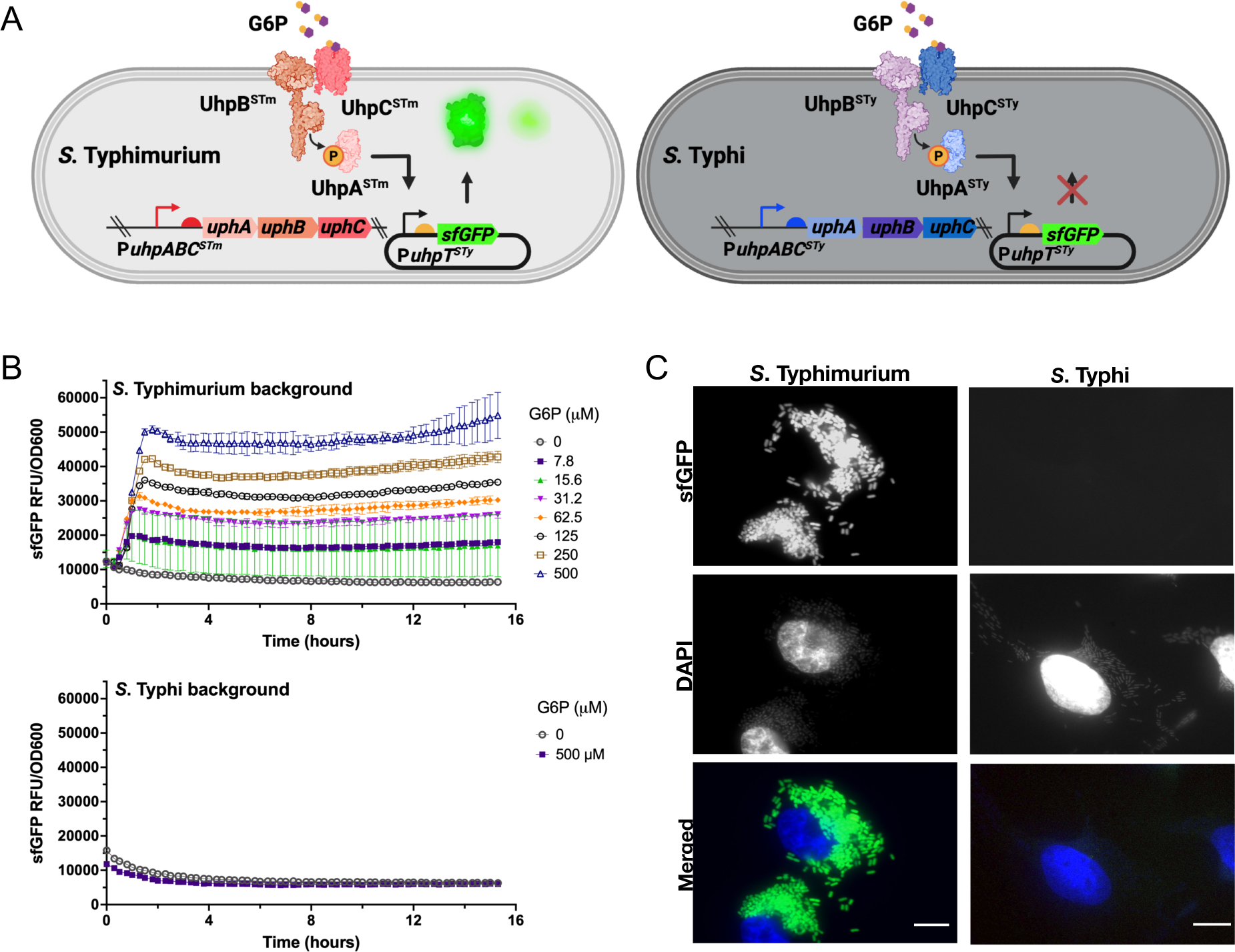
(**A**) Diagram of the G6P Biosensor for detection of glucose-6-phosphate in *S*. Typhimurium and *S*. Typhi. The membrane UhpC sensor detects G6P on the extracellular environment and interacts with the membrane UhpB histidine kinase that phosphorylates the cytosolic UhpA response regulator, which activates the P*uhpT* promoter. The *S.* Typhi P*uhpT* promoter was coupled to a *sfGFP* reporter gene on a plasmid system and transformed into *S*. Typhimurium and *S*. Typhi. In the presence of G6P, the cognate promoter P*uhpT^STy^* is activated leading to sfGFP expression. The possible steps where the sensing and regulation could be impaired in *S*. Typhi are noted. (**B**) Test of the response of the *S*. Typhimurium and *S*. Typhi G6P biosensors after growth in media containing increasing concentrations of G6P. The sfGFP fluorescense signal over time for the *S*. Typhimurium (upper graph) and *S*. Typhi (lower graph) G6P biosensors are shown. Values indicate the mean ± SEM of n= 3 replicates per condition. (**C**) Functionality of the G6P Biosensor in *S.* Typhimurium and *S.* Typhi in the context of cultured epithelial cell infection. Fluorescence microscopy of HeLa cells infected with bacterial strains encoding the G6P biosensors. HeLa cells were infected with wild type *S*. Typhimurium and *S*. Typhi strains encoding the G6P biosensors for 20 hs, fixed, stained with DAPI, and examined under a fluorescence microscope. For all images, brightness and contrast were adjusted for each of the individual channels to maximize visual clarity using the same parameters for both strains. Scale bars, 10 μm.

**Figure S2.**
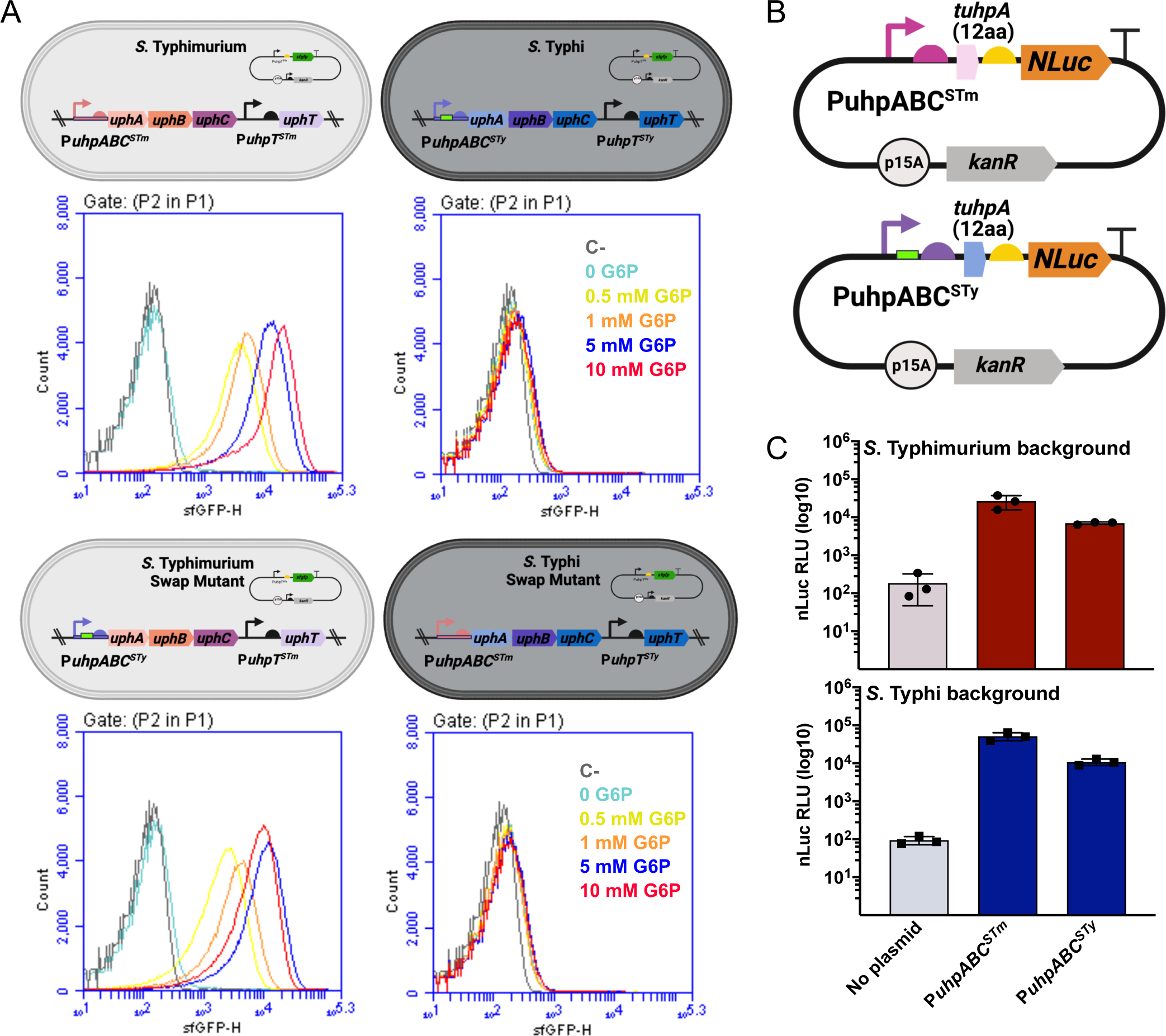
Investigation of the *S*. Typhi P*uhpABC* promoter. (**A**) Wild-type and mutant strains of *S*. Typhimurium containing the P*uhpABC* promoter from *S*. Typhi (P*uhpABC^STy^*), or *S*. Typhi containing the P*uhpABC* promoter from *S*. Typhimurium (P*uhpABC^STm^*), both harboring the G6P biosensor (P*uhpT*-sfGFP) were grown in presence of G6P for 20 hs and analyzed by flow-cytometry. Histograms show the sfGFP fluorescence intensities of individual bacteria for the indicated concentration of G6P. (**B**) Diagram of the transcriptional reporters in which the P*uhpABC^STm^*or the P*uhpABC^STy^* (containing an 80 bp insert indicated with a green rectangle) promoters drive the expression of a NanoLuc luciferase (NLuc). (**C**) *S*. Typhimurium and *S*. Typhi harboring the P*uhpABC^STm^*-*NLuc* or P*uhpABC^STy^*-*NLuc* reporters were grown for 20 hs, lysed and the luminescence was measured on a microplate reader. The signal in Relative Luminescence (RLU) for the indicated strains is shown. No statistically significative difference was observed between the P*uhpABC^STm^*-*NLuc* or P*uhpABC^STy^*-*NLuc* reporters as determined by Anova with Dunnett’s multiple comparisons test.

**Figure S3.**
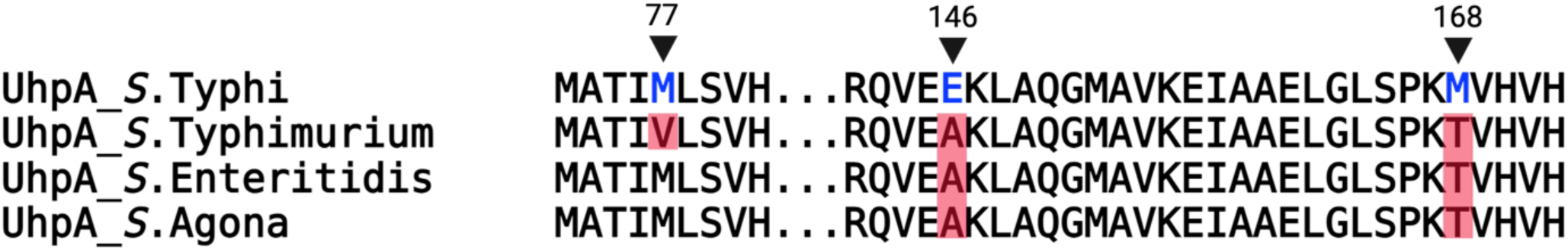
Alignment of portions of the amino acid sequence of the transcriptional regulator UhpA showing differences between *S*. Typhi CT18 and nontyphoidal *S. enterica* serovars *S.* Typhimurium LT2, *S.* Enteritidis P125109 and *S.* Agona SL483. The rest of the sequences not shown are identical between the different serovars.

**Figure S4.**
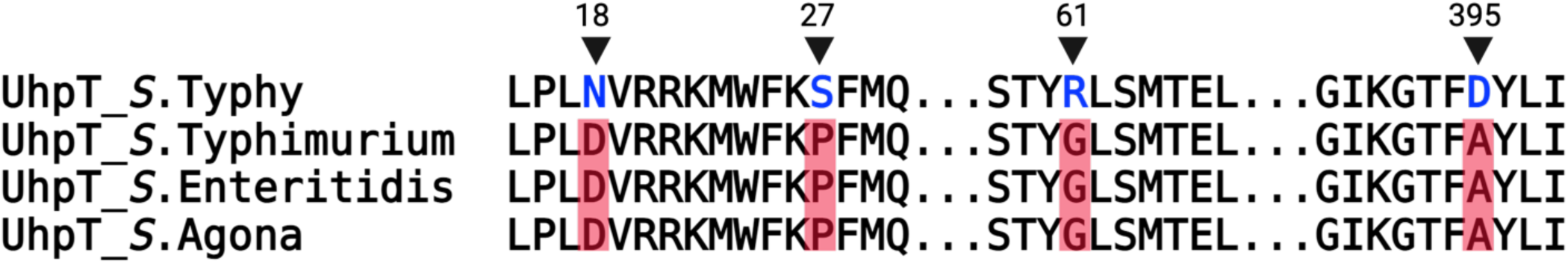
Alignment of portions of the amino acid sequence of the antiporter UhpT showing differences between *S*. Typhi CT18 and nontyphoidal *S. enterica* serovars *S.* Typhimurium LT2, *S.* Enteritidis P125109 and *S.* Agona SL483. The rest of the sequences not shown are identical between the different serovars.

**Figure S5.**
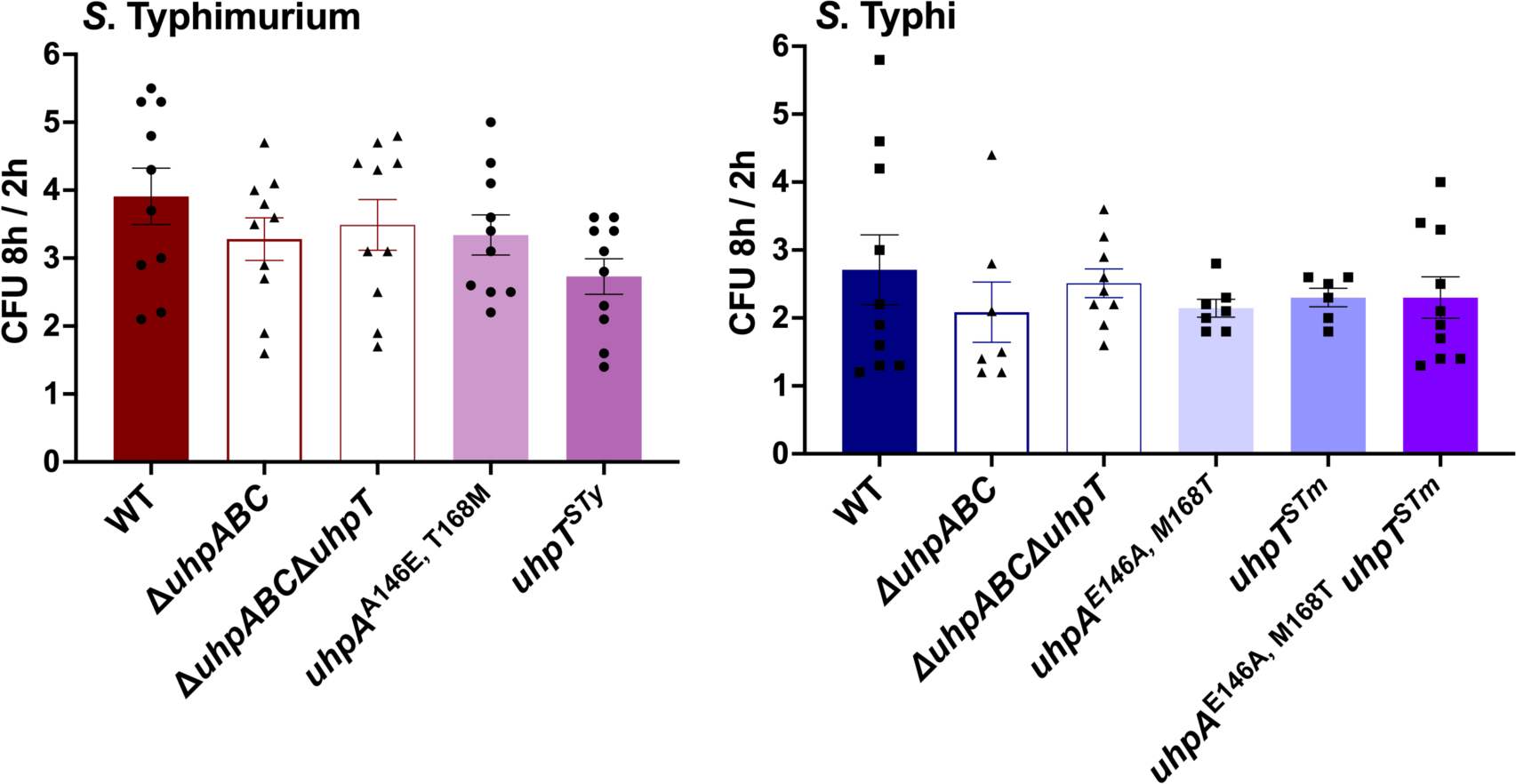
Intracellular growth of different *S*. Typhimurium or *S.* Typhi *uhp* mutants. Cultured HeLa cells were infected with indicated bacterial strains and the CFU 2 and 8 hours after infection were determined by plate dilutions. Values represent the ratio between the CFUs measured at 8 and 2 hours post infection and are the mean ± SEM of n= 7 - 10 replicates per condition. No statistically significative difference was observed as determined using Anova with Dunnett’s multiple comparisons test.

**Figure S6.**
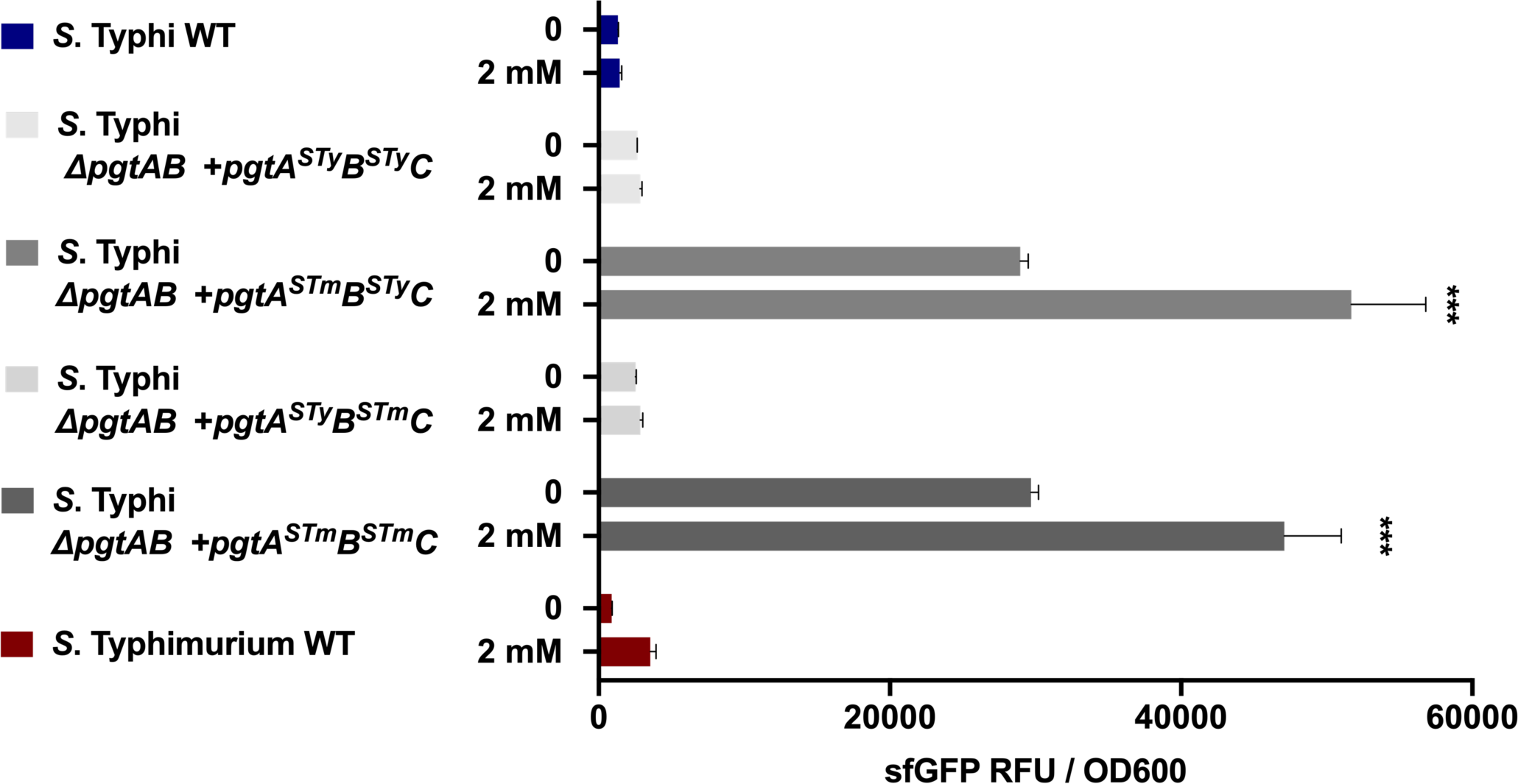
Investigation of the functionality of the PgtA and PgtB regulatory proteins. The *S.* Typhi *pgtA* and *pgtB* genes were swapped for their *S*. Typhimurium homologs in the 3PG biosensor plasmid. The resulting constructs were introduced into the *S*. Typhi *ΔpgtAB* mutant background and tested on the presence or absence of 3PG at 2 mM. The fluorescent gene expression normalized to cell density (sfGFP RFU / OD600) is shown. Values indicate the mean ± SEM of n= 3 replicates per condition. Asterisks denote statistically significant differences relative to the corresponding uninduced sample determined using Anova with Dunnett’s multiple comparisons test. ***p < 0.001.

**Figure S7.**
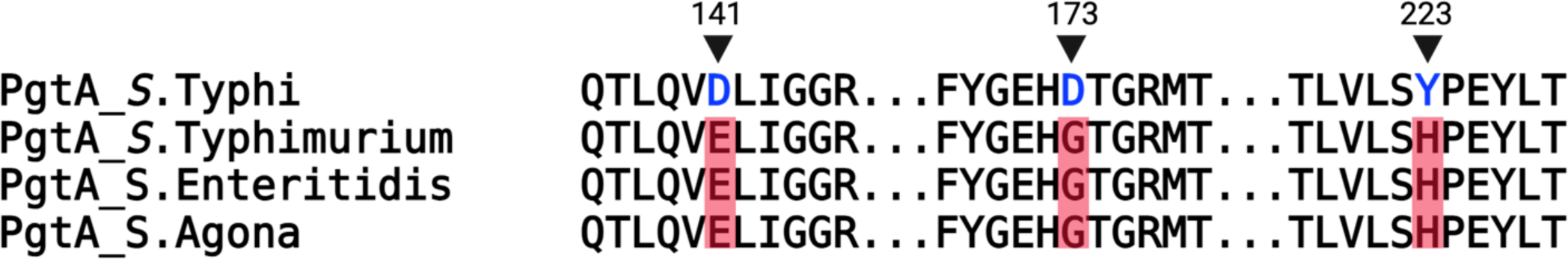
Alignment of portions of the amino acid sequence of the transcriptional regulator PgtA showing differences between *S*. Typhi CT18 and nontyphoidal *S. enterica* serovars *S.* Typhimurium LT2, *S.* Enteritidis P125109 and *S.* Agona SL483. The rest of the sequences not shown are identical between the different serovars.

**Figure S8.**
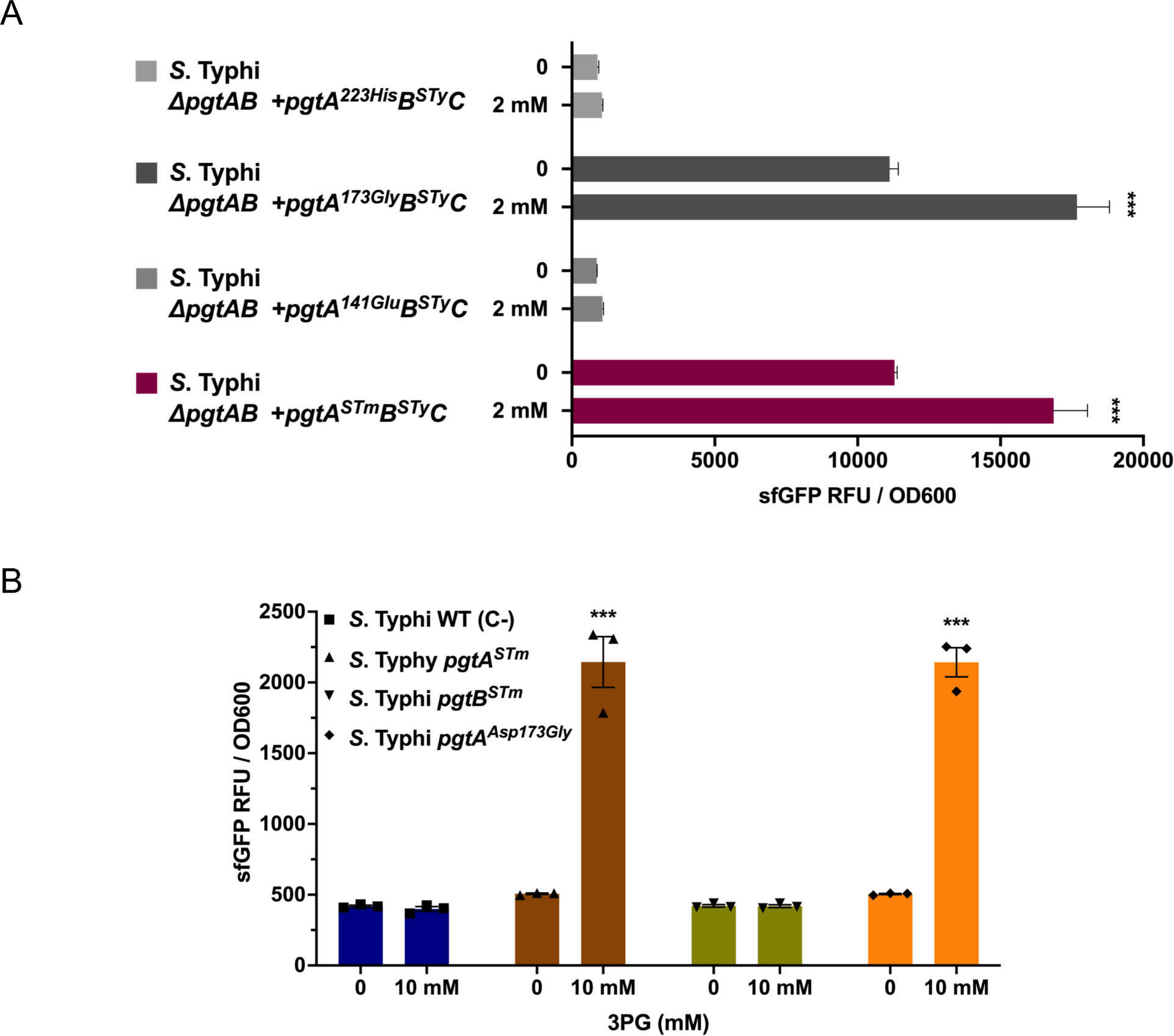
Investigation of the functionality of *S*. Typhi PgtA amino acid substitutions. (**A**) *S*. Typhi PgtA amino acids at the indicated positions were substituted for the equivalent *S*. Typhimurium residues in the plasmid backbone containing the 3PG biosensor. The resulting constructs were introduced into the *S*. Typhi *ΔpgtAB* mutant background and the transcriptional response was tested in the presence or absence of 3PG at 2mM. Values represent the fluorescence signal normalized to cell density (sfGFP RFU / OD600) and are the mean ± SEM of n= 4 replicates per condition. (**B**) The S. Typhi *pgtA* or *pgtB* were individually swapped for their S. Typhimurium homologues (S. Typhi *pgtA^Stm^* and S. Typhi *pgtB^Stm^).* The resulting strains and an *S*. Typhi strain encoding the *pgtA^Asp173Gly^,* all encoding the 3PG biosensor, were tested for their transcriptional response to 3PG at 10 mM. The fluorescence signal normalized to cell density (sfGFP RFU / OD600) is shown. Values indicate the mean ± SEM of n= 3 replicates per condition. For all panels, asterisks denote statistically significant differences relative to the corresponding uninduced sample determined using Anova with Dunnett’s multiple comparisons test. ***p < 0.001.

**Figure S9.**
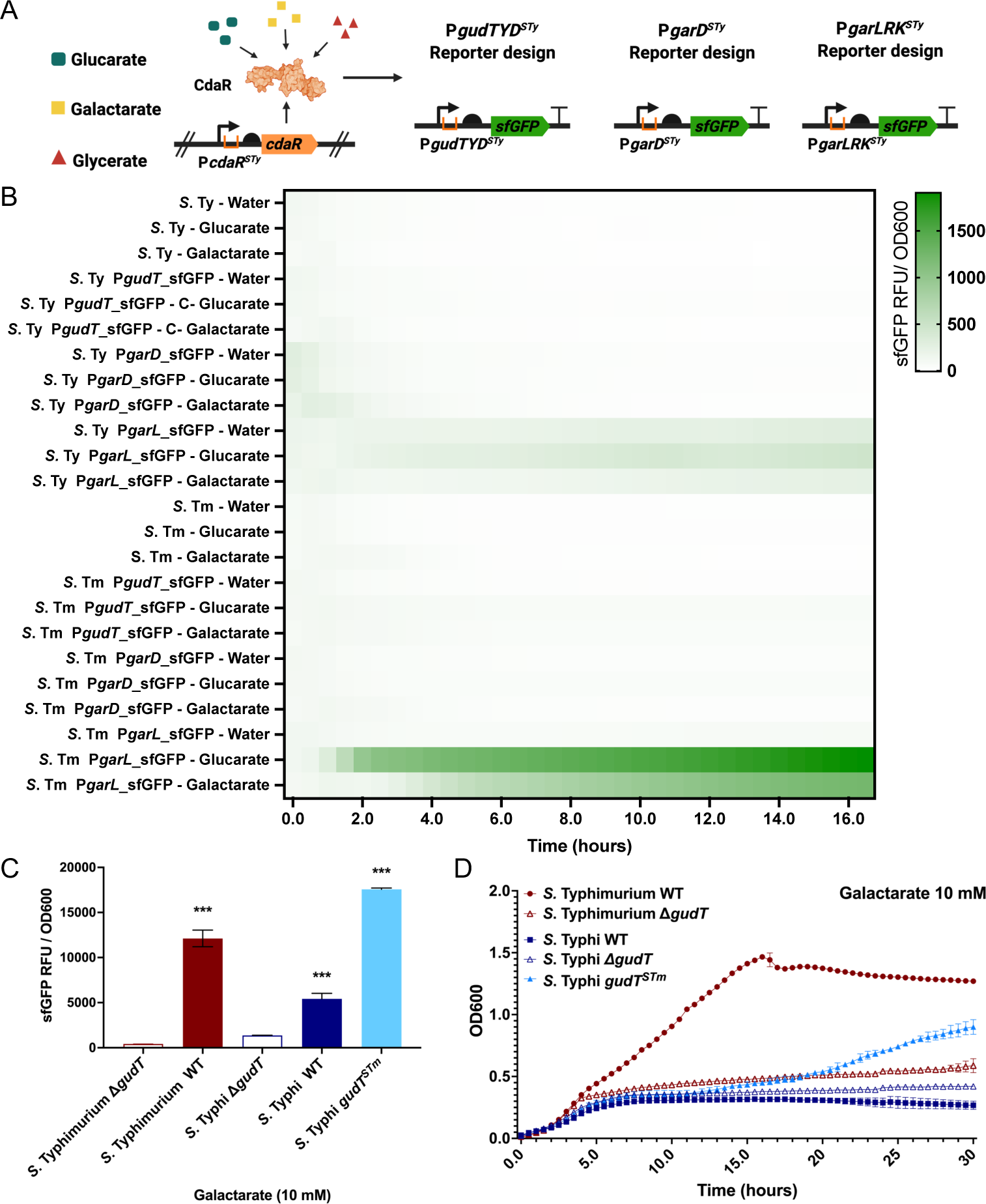
(**A**) (**A**) Diagram of the biosensor for the detection of glucarate, galactarate and glycerate based on the CdaR transcription factor and the cognate promoters P*gudTYD*, P*garD* and P*garLRK* driving expression of *sfGFP*. (**B**) The indicated wild-type strains of *S*. Typhimurium (S. Tm) and *S*. Typhi (S. Ty) harboring the indicated biosensors were tested in LB medium containing glucarate or galactarate (1 mM), as indicated, or water as a negative control, and the fluorescence signal (normalized to cell density) was measured (sfGFP RFU/OD600) for each strain. (**C**) The indicated *S*. Typhimurium and *S*. Typhi mutant strains all harboring the P*garL*_*sfGFP* transcriptional reporter were grown in LB medium containing galactarate at 10mM, and the fluorescence signal (normalized to cell density) was measured (sfGFP RFU/OD600) for each strain. Values are the mean ± SEM of n= 3 replicates per condition. Asterisks denote statistically significant differences determined using Anova with Dunnett’s multiple comparisons test. ***p < 0.001. (**D**) Growth kinetics of wild-type *S.* Typhimurium and *S.* Typhi or their isogenic mutants *ΔgudT, ΔgudT* and *gudT^STm^* as indicated. Growth was monitored in M9 +0.05% casamino acids medium containing 10 mM galactarate as the only carbon source. The OD600 was measured at 30 min intervals and values represent the mean ± SEM of n=3-4 replicates per condition.

**Figure S10.**
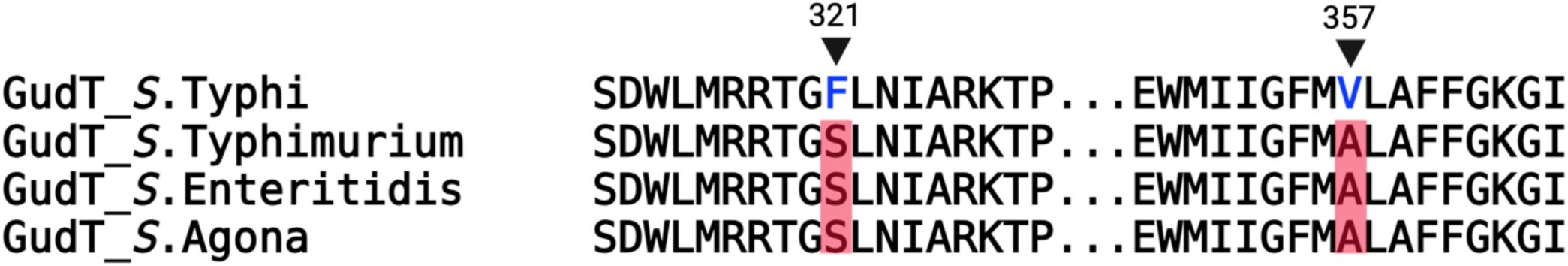
Alignment of portions of the amino acid sequence of the glucarate/galactarate GudT transporter showing differences between *S*. Typhi CT18 and nontyphoidal *S. enterica* serovars *S.* Typhimurium LT2, *S.* Enteritidis P125109 and *S.* Agona SL483. The rest of the sequences not shown are identical between the different serovars.

**Figure S11.**
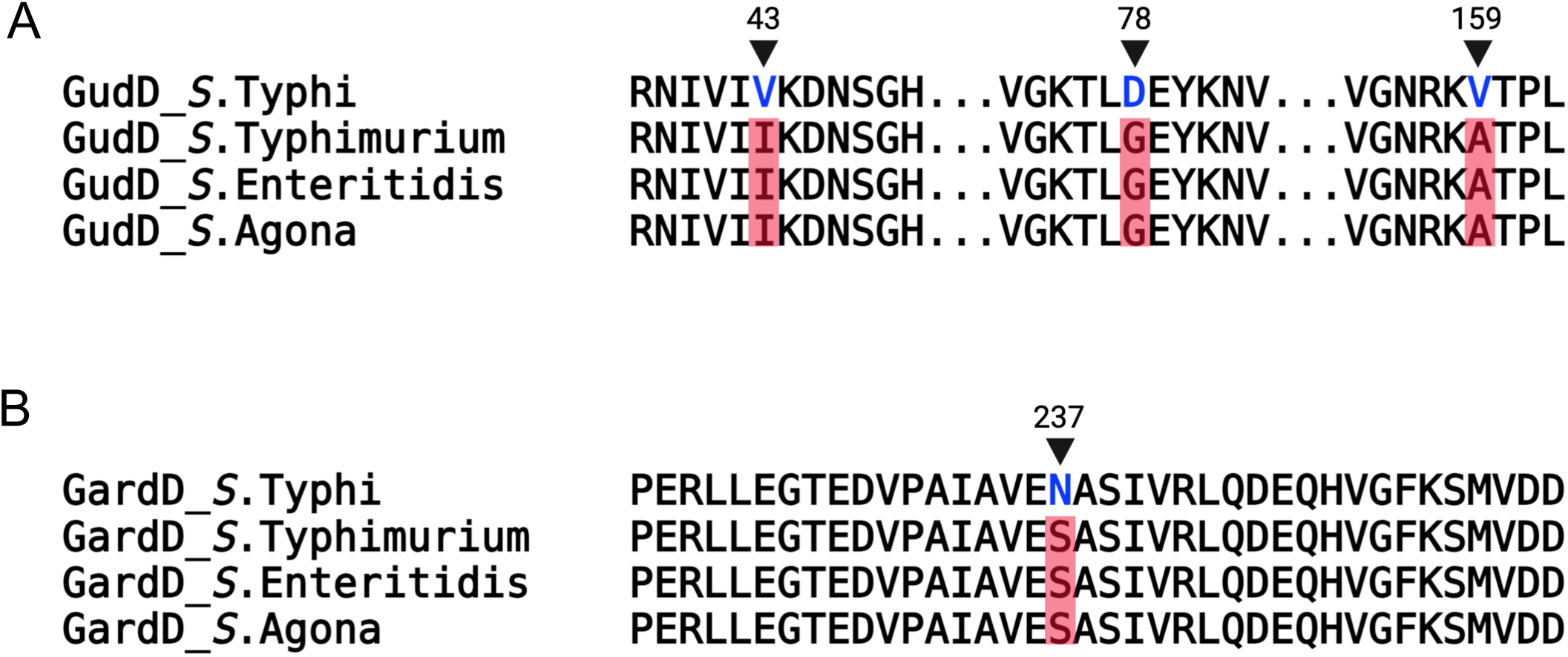
Alignment of portions of the amino acid sequences of the GudD (A) and GarD (B) showing differences between *S*. Typhi CT18 and nontyphoidal *S. enterica* serovars *S.* Typhimurium LT2, *S.* Enteritidis P125109 and *S.* Agona SL483. The rest of the sequences not shown are identical between the different serovars.

**Figure S12.**
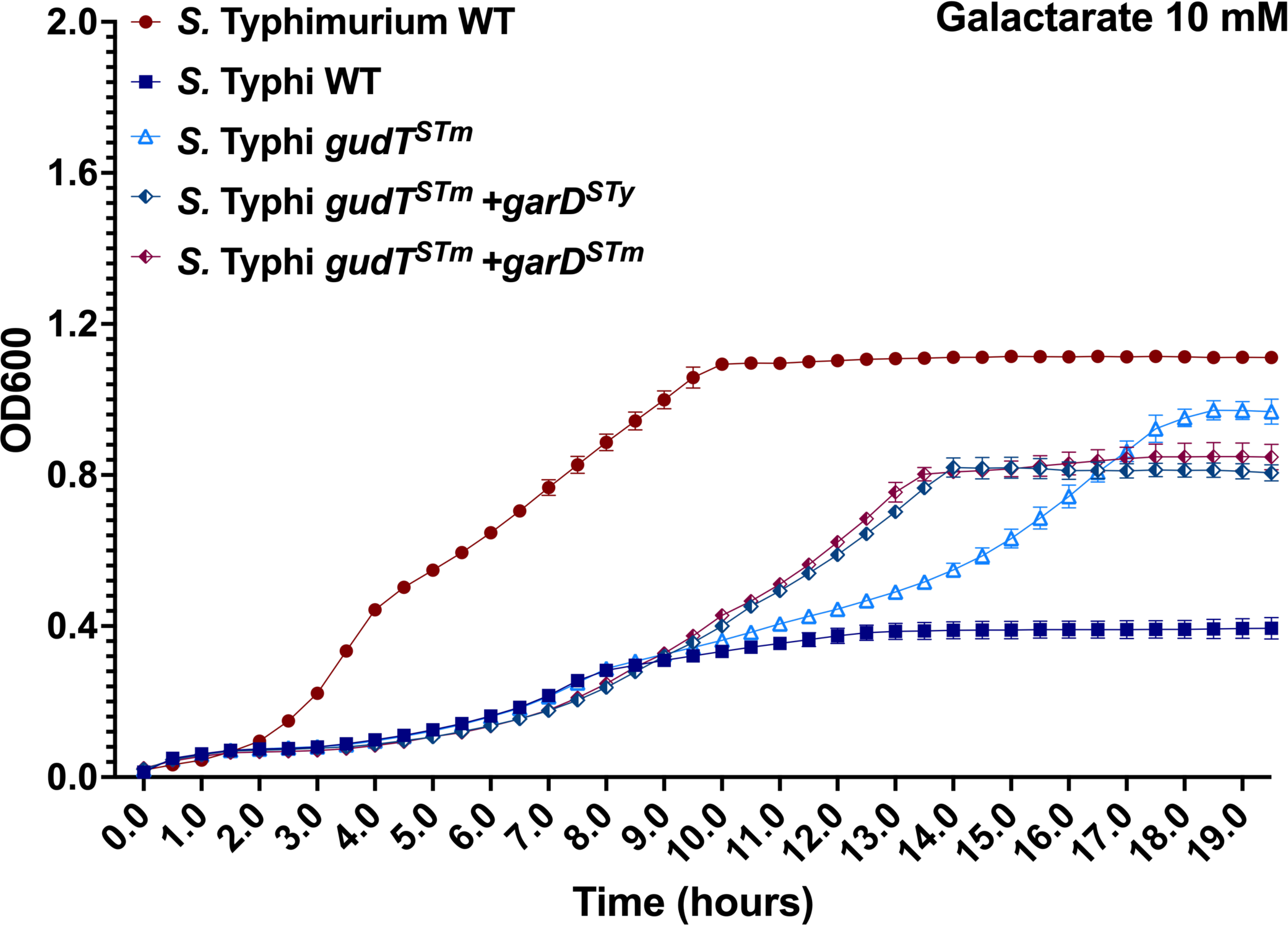
Growth kinetics of *S.* Typhi strains expressing *S*. Typhimurium *gudT* (*gudT^STm^*) and *garD* (*garD^STm^*) or (*garD^STy^*) alleles as indicated. Strains were grown in M9 +0.05% casamino acids medium containing 10 mM galactarate as the only carbon source, and the OD600 was measured at 30 min intervals. Values indicate the mean ± SEM of n= 3-4 replicates per condition.

**Fig S13.**
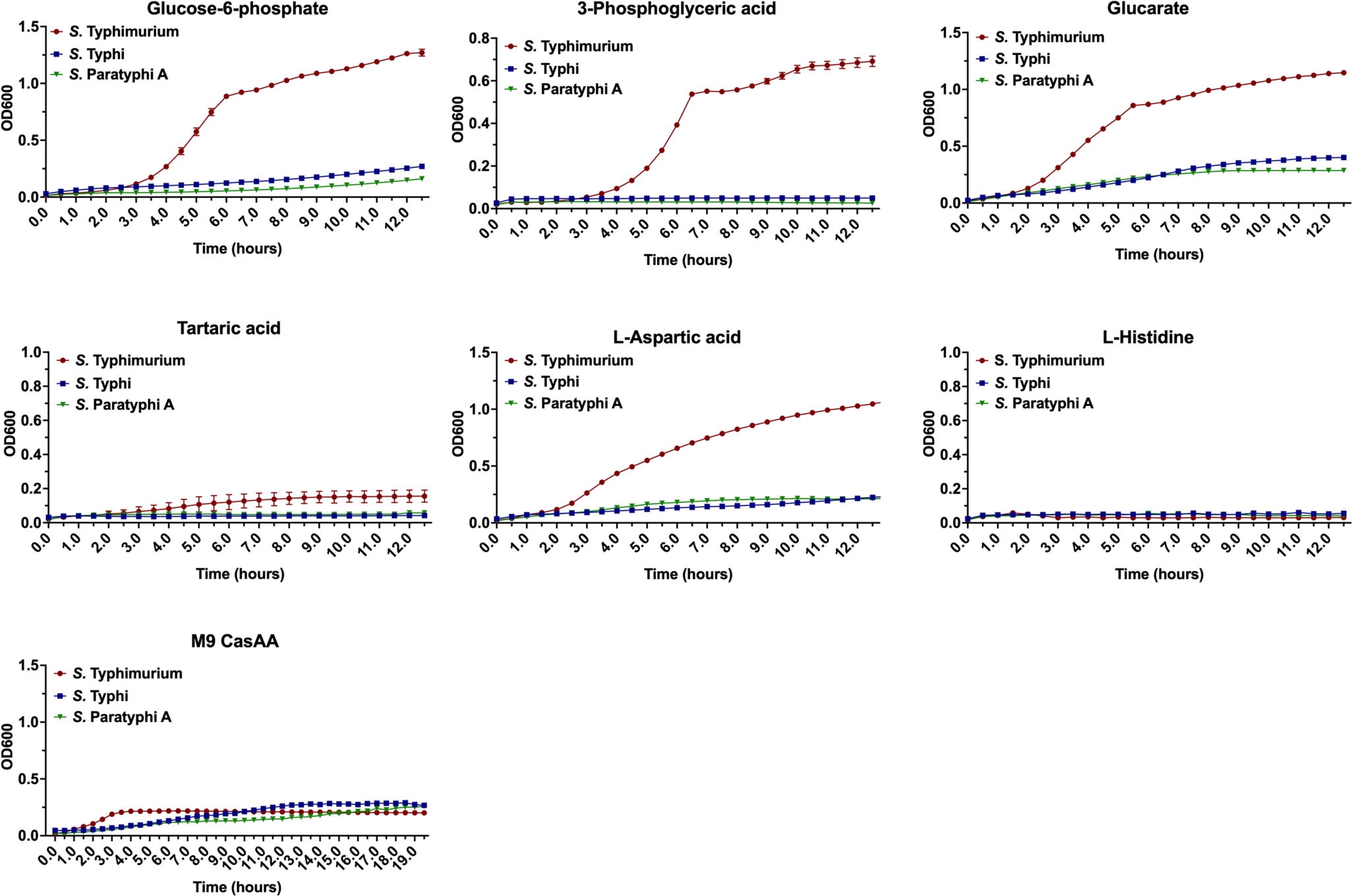
Growth kinetics of *S*. Typhimurium, *S*. Typhi and *S*. Paratyphi A in minimal medium containing glucose-6-phosphate, 3-phosphoglyceric acid, glucarate, tartrate, aspartate, or L-histidine as the sole carbon source. Bacteria were grown in M9 medium supplemented with casamino acids (0.05%) and containing 20 mM of the different metabolites and the OD600 was measured at 30 min intervals. Values indicate the mean ± SEM of n= 3-4 replicates per condition.

**Figure S14.**
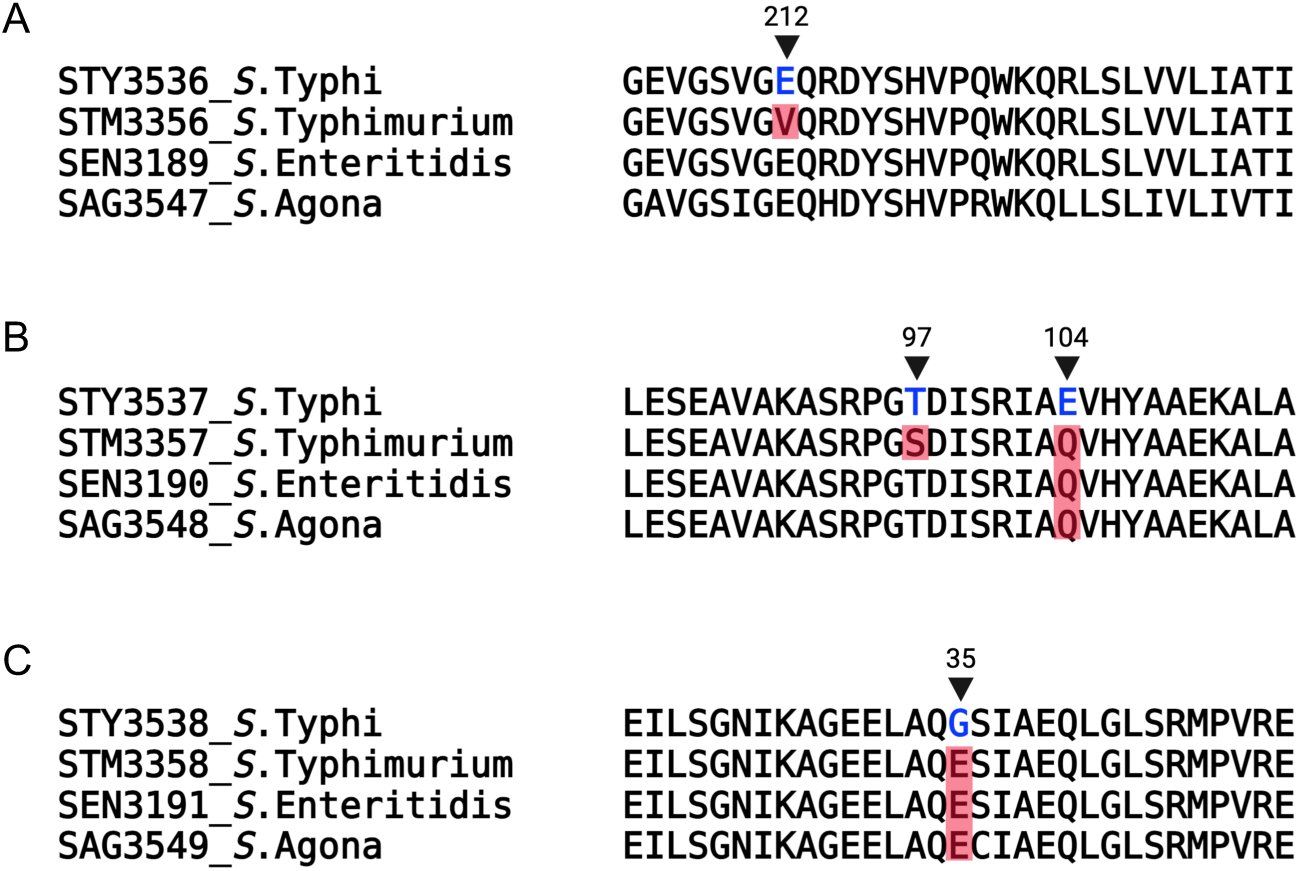
Alignment of portions of the amino acid sequence of *S*. Typhi STY3536 (**A**), STY3537 (**B**), and STY3538 (**C**) showing differences between *S*. Typhi CT18 and nontyphoidal *S. enterica* serovars *S.* Typhimurium LT2, *S.* Enteritidis P125109 and *S.* Agona SL483. The rest of the sequences not shown are identical between the different serovars.

**Figure S15.**
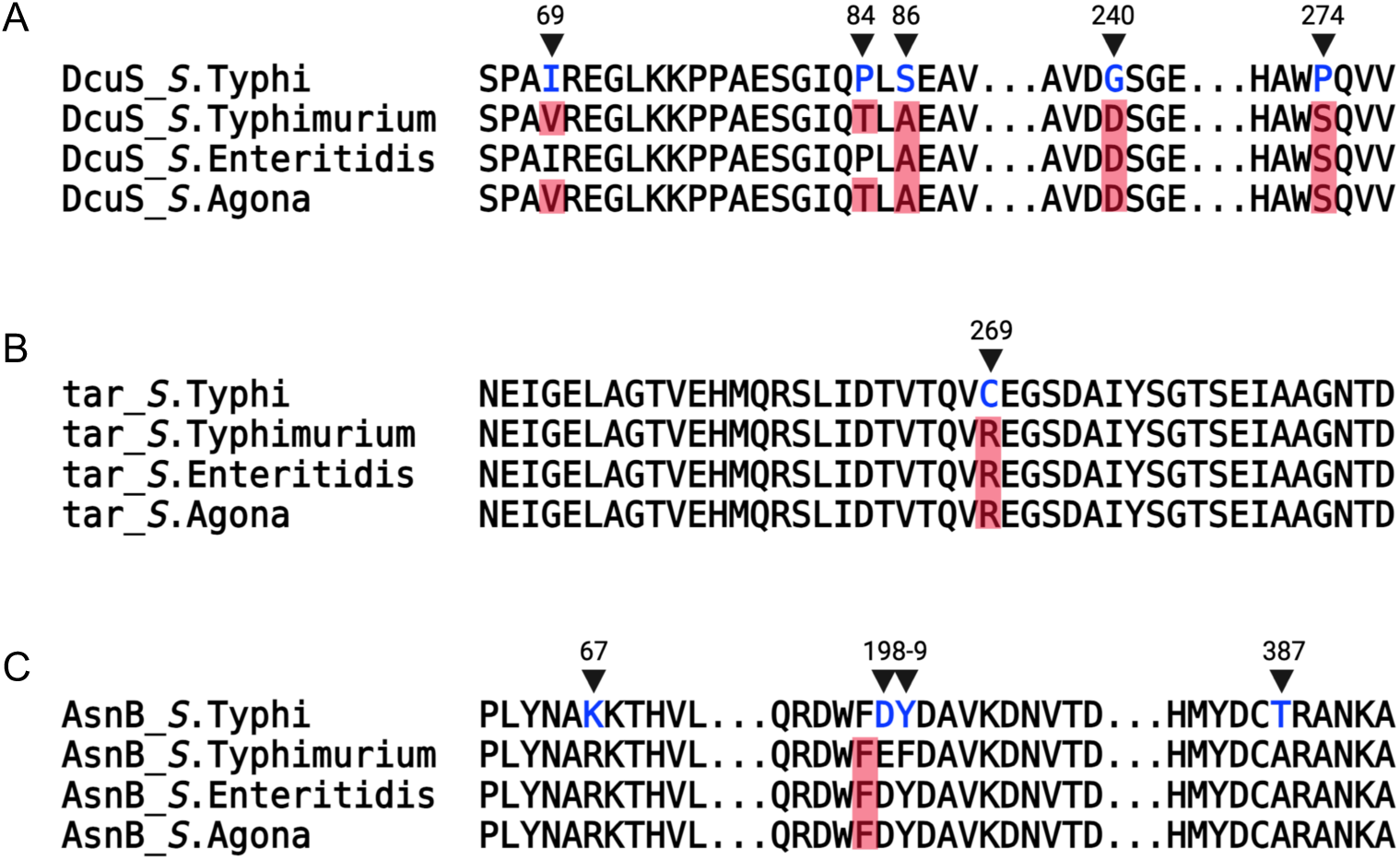
Alignment of portions of the amino acid sequence of *S*. Typhi DcuS (**A**), Tar (**B**), and AsnB (**C**) showing differences between *S*. Typhi CT18 and nontyphoidal *S. enterica* serovars *S.* Typhimurium LT2, *S.* Enteritidis P125109 and *S.* Agona SL483. The rest of the sequences not shown are identical between the different serovars.

**Figure S16.**
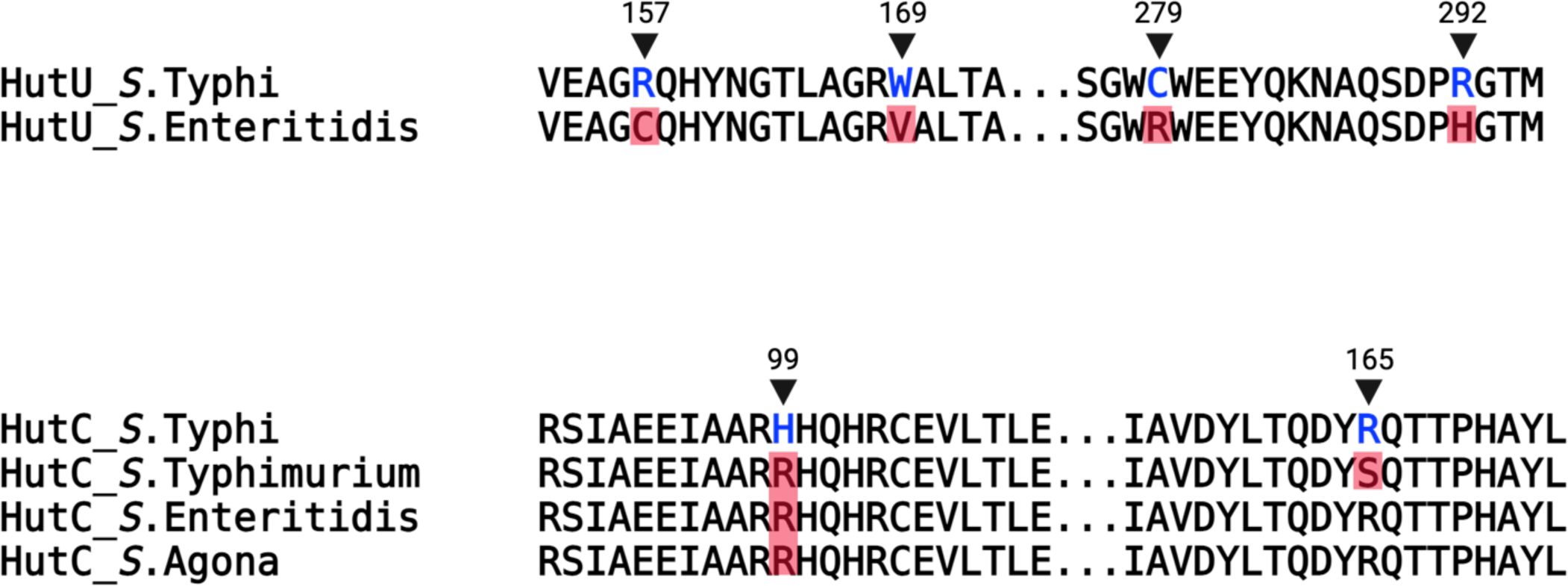
Alignment of portions of the amino acid sequence of *S*. Typhi HutU (**A**) and HutC (**B**) showing differences between *S*. Typhi CT18 and nontyphoidal *S. enterica* serovars *S.* Typhimurium LT2, *S.* Enteritidis P125109 and *S.* Agona SL483. The rest of the sequences not shown are identical between the different serovars.

## Notes

### Competing Interest Statement

The authors have declared no competing interest.

## References

1. Collaborators. GN-TSID. 2019. The global burden of non-typhoidal salmonella invasive disease: a systematic analysis for the Global Burden of Disease Study 2017. Lancet Infect Dis 19:1312–1324.

2. Balasubramanian R, Im J, Lee J, Jeon H, Mogeni O, Kim J, Rakotozandrindrainy R, Baker S, Marks F. 2019. The global burden and epidemiology of invasive non-typhoidal Salmonella infections. Hum Vaccin Immunother 15:1421–1426.

3. Mogasale V, Maskery B, Ochiai RL, Lee JS, Mogasale VV, Ramani E, Kim YE, Park JK, Wierzba TF. 2014. Burden of typhoid fever in low-income and middle-income countries: a systematic, literature-based update with risk-factor adjustment. Lancet Glob Health 2:e570–80.

4. Shannon E, Majowicz S, Musto J, Scallan E, Angulo F, Kirk M, O’Brien S, Jones T, Fazil A, Hoekstra R. 2010. The Global Burden of Nontyphoidal Salmonella Gastroenteritis. Clinical Infectious Diseases 50,:882–889.

5. Brenner F, Villar R, Angulo F, Tauxe R, Swaminathan B. 2000. Salmonella nomenclature. J Clin Microbiol 38:2465–7.

6. Dougan G, Baker S. 2014. Salmonella enterica serovar Typhi and the pathogenesis of typhoid fever. Annu Rev Microbiol 68:317–36.

7. Ohl ME, Miller SI. 2001. Salmonella: a model for bacterial pathogenesis. Annu Rev Med 52:259–274.

8. Raffatellu M, Wilson R, Winter S, Bäumler A. 2008. Clinical pathogenesis of typhoid fever. J Infect Dev Ctries 2:260–6.

9. Grassl G, Finlay B. 2008. Pathogenesis of enteric Salmonella infections. Curr Opin Gastroenterol 24:22–6

10. Parry C, Hien TT, Dougan G, White N, Farrar J. 2002. Typhoid fever. N Engl J Med 347:1770–82.

11. Buckle G, Walker C, Black R. 2012. Typhoid fever and paratyphoid fever: Systematic review to estimate global morbidity and mortality for 2010. J Glob Health 2:010401.

12. Kim JH, Mogasale V, Im J, Ramani E, Marks F. 2017. Updated estimates of typhoid fever burden in sub-Saharan Africa. Lancet Glob Health 5:e969.

13. McClelland M, Sanderson KE, Clifton SW, Latreille P, Porwollik S, Sabo A, Meyer R, Bieri T, Ozersky P, McLellan M, Harkins CR, Wang C, Nguyen C, Berghoff A, Elliott G, Kohlberg S, Strong C, Du F, Carter J, Kremizki C, Layman D, Leonard S, Sun H, Fulton L, Nash W, Miner T, Minx P, Delehaunty K, Fronick C, Magrini V, Nhan M, Warren W, Florea L, Spieth J, Wilson RK. 2004. Comparison of genome degradation in Paratyphi A and Typhi, human-restricted serovars of Salmonella enterica that cause typhoid. Nat Genet 36:1268–74.

14. Holt K, Thomson N, Wain J, Langridge G, Hasan R, Bhutta Z, Quail M, Norbertczak H, Walker D, Simmonds M, White B, Bason N, Mungall K, Dougan G, Parkhill J. 2009. Pseudogene accumulation in the evolutionary histories of Salmonella enterica serovars Paratyphi A and Typhi. BMC Genomics 10:36.

15. Didelot X, Achtman M, Parkhill J, Thomson NR, Falush D. 2007. A bimodal pattern of relatedness between the Salmonella Paratyphi A and Typhi genomes: Convergence or divergence by homologous recombination? Genome Research 17:61–61.

16. Langridge G, Fookes M, Connor T, Feltwell T, Feasey N, BN P, Seth-Smith H, Barquist L, Stedman A, Humphrey T, Wigley P, Peters S, Maskell D, Corander J, Chabalgoity J, Barrow P, Parkhill J, Dougan G, Thomson N. 2015. Patterns of genome evolution that have accompanied host adaptation in Salmonella. Proc Natl Acad Sci U S A 112:Patterns of genome evolution that have accompanied host adaptation in Salmonella. Proc Natl Acad Sci U S A. 2015 Jan 20;112(3):863–8. doi: 10.1073/pnas.1416707112. Epub 2014 Dec 22. PMID: 25535353; PMCID: PMC4311825.

17. Nuccio S, Bäumler A. 2014. Comparative analysis of Salmonella genomes identifies a metabolic network for escalating growth in the inflamed gut. mBio 5:e00929–14.

18. Manesh A, Meltzer E, Jin C, Britto C, Deodhar D, Radha S, Schwartz E, Rupali P. 2021. Typhoid and paratyphoid fever: a clinical seminar. J Travel Med 28:taab012.

19. Parkhill J, Dougan G, James KD, Thomson NR, Pickard D, Wain J, Churcher C, Mungall KL, Bentley SD, Holden MT, Sebaihia M, Baker S, Basham D, Brooks K, Chillingworth T, Connerton P, Cronin A, Davis P, Davies RM, Dowd L, White N, Farrar J, Feltwell T, Hamlin N, Haque A, Hien TT, Holroyd S, Jagels K, Krogh A, Larsen TS, Leather S, Moule S, O’Gaora P, Parry C, Quail M, Rutherford K, Simmonds M, Skelton J, Stevens K, Whitehead S, Barrell BG. 2001. Complete genome sequence of a multiple drug resistant Salmonella enterica serovar Typhi CT18. Nature 413:848–52.

20. Runyen-Janecky L, Payne S. 2002. Identification of chromosomal Shigella flexneri genes induced by the eukaryotic intracellular environment. Infect Immun 70:4379–88.

21. Powers T, Haeberle A, Predeus A, Hammarlöf D, Cundiff J, Saldaña-Ahuactzi Z, Hokamp K, Hinton J, Knodler L. 2021. Intracellular niche-specific profiling reveals transcriptional adaptations required for the cytosolic lifestyle of Salmonella enterica. PLoS Pathog 17:e1009280.

22. Knodler L. 2015. Salmonella enterica: living a double life in epithelial cells. Curr Opin Microbiol 23:23–31.

23. Kadner R, Webber C, Island M. 1993. The family of organo-phosphate transport proteins includes a transmembrane regulatory protein. J Bioenerg Biomembr 25:637–45.

24. Island MD, Wei BY, Kadner RJ. 1992. Structure and function of the uhp genes for the sugar phosphate transport system in Escherichia coli and Salmonella typhimurium. Journal of Bacteriology 174:2754–2762.

25. Spinnenhirn V, Farhan H, Basler M, Aichem A, Canaan A, Groettrup M. 2014. The ubiquitin-like modifier FAT10 decorates autophagy-targeted Salmonella and contributes to Salmonella resistance in mice. Journal of Cell Science 127:4883–4893.

26. Finn CE, Chong A, Cooper KG, Starr T, Steele-Mortimer O. 2017. A second wave of Salmonella T3SS1 activity prolongs the lifespan of infected epithelial cells. PLOS Pathogens 13:e1006354–e1006354.

27. Chong A, Cooper KG, Kari L, Nilsson OR, Hillman C, Fleming BA, Wang Q, Nair V, Steele-Mortimer O. 2021. Cytosolic replication in epithelial cells fuels intestinal expansion and chronic fecal shedding of Salmonella Typhimurium. Cell Host & Microbe doi:10.1016/j.chom.2021.04.017.

28. DeBerardinis R, Lum J, Hatzivassiliou G, Thompson C. 2008. The biology of cancer: metabolic reprogramming fuels cell growth and proliferation. Cell Metab 7:11–20.

29. Palsson-McDermott E, O’Neill L. 2013. The Warburg effect then and now: from cancer to inflammatory diseases. Bioessays. Bioessays 35:965–73.

30. Escoll P, Buchrieser C. 2018. Metabolic reprogramming of host cells upon bacterial infection: Why shift to a Warburg-like metabolism? The FEBS Journal 285:2146–2160.

31. Jiang L, Wang P, Song X, Zhang H, Ma S, Wang J, Li W, Lv R, Liu X, Ma S, Yan J, Zhou H, Huang D, Cheng Z, Yang C, Feng L, Wang L. 2021. Salmonella Typhimurium reprograms macrophage metabolism via T3SS effector SopE2 to promote intracellular replication and virulence. Nature Communications 2021 12:1 12:1–18.

32. Jiang S-Q, Yu G-Q, Li Z-G, Hong J-S. 1988. Genetic Evidence for Modulation of the Activator by Two Regulatory Proteins Involved in the Exogenous Induction of Phosphoglycerate Transport in Salmonella typhimurium. JOURNAL OF BACTERIOLOGY 170:4304–4308.

33. Niu S, Jiang SQ, Hong J. 1995. Salmonella typhimurium pgtB mutants conferring constitutive expression of phosphoglycerate transporter pgtP independent of pgtC. Journal of Bacteriology 177:4297–4302.

34. Faber F, Tran L, Byndloss MX, Lopez CA, Velazquez EM, Kerrinnes T, Nuccio S-P, Wangdi T, Fiehn O, Tsolis RM, Bäumler AJ. 2016. Host-mediated sugar oxidation promotes post-antibiotic pathogen expansion. Nature 534:697–699.

35. Monterrubio R, Baldoma L, Obradors N, Aguilar J, Badia J. 2000. A common regulator for the operons encoding the enzymes involved in D-galactarate, D-glucarate, and D-glycerate utilization in Escherichia coli. Journal of Bacteriology 182:2672–2674.

36. Lamichhane-Khadka R, Frye JG, Porwollik S, McClelland M, Maier RJ. 2011. Hydrogen-stimulated carbon acquisition and conservation in Salmonella enterica serovar Typhimurium. Journal of bacteriology 193:5824–5832.

37. Zientz E BJ, Unden G.. 1998. Fumarate regulation of gene expression in Escherichia coli by the DcuSR (dcuSR genes) two-component regulatory system. J Bacteriol 180:5421–5.

38. Felton J, Michaelis S, Wright A. 1980. Mutations in two unlinked genes are required to produce asparagine auxotrophy in Escherichia coli. J Bacteriol 142:221–8.

39. Smith G, Magasanik B. 1971. The two operons of the histidine utilization system in Salmonella typhimurium. J Biol Chem 246:3330–41.

40. Hoiseth SK, Stocker BA. 1981. Aromatic-dependent *Salmonella typhimurium* are non-virulent and effective as live vaccines. Nature 291:238–239.

41. Galan JE, Curtiss R, 3rd. 1991. Distribution of the invA, -B, -C, and -D genes of Salmonella typhimurium among other Salmonella serovars: invA mutants of Salmonella typhi are deficient for entry into mammalian cells. Infect Immun 59:2901–8.

42. Penfold R, Pemberton J. 1992. An improved suicide vector for construction of chromosomal insertion mutations in bacteria. Gene 118:145–6.

43. Demarre G, Guerout AM, Matsumoto-Mashimo C, Rowe-Magnus DA, Marliere P, Mazel D. 2005. A new family of mobilizable suicide plasmids based on broad host range R388 plasmid (IncW) and RP4 plasmid (IncPα) conjugative machineries and their cognate Escherichia coli host strains. Res Microbiol 156:245–255.

44. Gibson D, Young L, Chuang R, Venter J, Hutchison Cr, Smith H. 2009. Enzymatic assembly of DNA molecules up to several hundred kilobases. Nat Methods 6:343–5

45. Meyer AJ, Segall-Shapiro TH, Glassey E, Zhang J, Voigt CA. 2019. Escherichia coli “Marionette” strains with 12 highly optimized small-molecule sensors. Nature Chemical Biology 15:196–204.

46. Kaniga K, Bossio JC, Galan JE. 1994. The Salmonella typhimurium invasion genes invF and invG encode homologues of the AraC and PulD family of proteins. Mol Microbiol 13:555–68.

47. Rogers JK, Guzman CD, Taylor ND, Raman S, Anderson K, Church GM. 2015. Synthetic biosensors for precise gene control and real-time monitoring of metabolites. Nucleic Acids Research 43:7648–7660.

48. Galán JE, Curtiss III R. 1990. Expression of *Salmonella typhimurium* genes required for invasion is regulated by changes in DNA supercoiling. Infect Immun 58:1879–1885.

49. Mirdita M, Schütze K, Moriwaki Y, Heo L, Ovchinnikov S, Steinegger M. 2022. ColabFold: making protein folding accessible to all. Nature Methods 19:679–682.

50. DeLano WL. 2002. The PyMOL Molecular Graphics System. http://www.pymolorg

